# Multivalent Human Cytomegalovirus Glycoprotein B Nucleoside-Modified mRNA Vaccines Demonstrate a Greater Breadth in T cell but not Antibody Responses

**DOI:** 10.1101/2022.11.23.517695

**Authors:** Hsuan-Yuan (Sherry) Wang, Leike Li, Cody S. Nelson, Richard Barfield, Sarah Valencia, Cliburn Chan, Hiromi Muramatsu, Paulo J.C. Lin, Norbert Pardi, Zhiqiang An, Drew Weissman, Sallie R. Permar

## Abstract

Human cytomegalovirus (HCMV) remains the most common congenital infection and infectious complication in immunocompromised patients. The most successful HCMV vaccine to-date, an HCMV glycoprotein B (gB) subunit vaccine adjuvanted with MF59, achieved 50% efficacy against primary HCMV infection. A previous study demonstrated that gB/MF59 vaccinees were less frequently infected with HCMV gB genotype strains most similar to the vaccine strain than strains encoding genetically distinct gB genotypes, suggesting strain-specific immunity accounted for the limited efficacy. To determine whether vaccination with multiple HCMV gB genotypes could increase the breadth of anti-HCMV gB humoral and cellular responses, we immunized 18 female rabbits with monovalent (gB-1), bivalent (gB-1+gB-3), or pentavalent (gB-1+gB-2+gB-3+gB-4+gB-5) gB lipid nanoparticle-encapsulated nucleoside-modified RNA (mRNA-LNP) vaccines. The multivalent vaccine groups did not demonstrate higher magnitude or breadth of the IgG response to the gB ectodomain or cell-associated gB compared to that of monovalent vaccine. Also, the multivalent vaccines did not show an increase in the breadth of neutralization activity and antibody-dependent cellular phagocytosis against HCMV strains encoding distinct gB genotypes. Yet, peripheral blood mononuclear cell-derived T cell responses elicited by multivalent vaccines were of a higher magnitude compared to that of monovalent vaccinated animals against a vaccine-mismatched gB genotype at peak immunogenicity. Our data suggests that inclusion of multivalent gB antigens is beneficial to increase the magnitude of T cell response but not an effective strategy to increase the breadth of anti-HCMV gB antibody responses. Further studies are required to validate whether the multivalent gB mRNA vaccines could effectively increase the T cell response breadth.

## Introduction

Human cytomegalovirus (HCMV), a ubiquitous β-herpesvirus, is a common cause of mild to severe disease in immunocompromised hosts and infants born with congenital infection ^1, 2^. The frequent and severe impacts of this infection has led to a HCMV vaccine being listed as a Tier 1 priority vaccine by the National Academy of Medicine for over 20 years ^3^. The dsDNA genome of HCMV is approximately 235 kb long and can encode up to 65 types of unique glycoproteins ^4^. HCMV glycoprotein B (gB), a relatively conserved membrane glycoprotein essential for viral entry into all host cells, has been identified as a vaccine target since 1980s due to its ability to elicit both neutralizing and non-neutralizing antibody responses ^5–9^. Previously, a gB subunit vaccine combined with the squalene adjuvant MF59, demonstrated modest effectiveness against virus acquisition in clinical trials ^10^. The gB/MF59 vaccine was administered to healthy HCMV-seronegative female adolescents and postpartum women and achieved approximately 50% efficacy against primary HCMV infection in phase II clinical trials ^11, 12^. In another phase II clinical trial, the vaccine-elicited gB-specific antibody titer was correlated with the reduction of post-transplant viremia in transplant recipients ^13^.Yet, the efficacy achieved with the HCMV gB-based vaccine will need to be enhanced in future vaccine products for clinical translation.

An HCMV vaccine design that incorporates the high degree of variation among HCMV strains is a possible strategy to improve vaccine efficacy through the enhancement of immunogenicity breadth. Infection with multiple HCMV strains, often referred to as mixed infection, is commonly observed in immunocompromised patients, such as AIDS patients ^14^ and organ transplant recipients ^15, 16^, including viruses with multiple gB genotypes in the same host. Mixed infection was also observed in infants with congenital HCMV infection, although the correlation between a specific HCMV genotype and clinical disease remained elusive ^17^. Besides acquisition of a new HCMV strain (known as re-infection), the replacement of one genotype for another genotype was observed ^18^. In immunocompromised patients, a new HCMV strain could also appear by reactivation and recombination with an existing HCMV strain ^19^. These studies all demonstrate that the diversity of HCMV strains may need to be considered in HCMV vaccine design.

HCMV gB, composed of ∼900 amino acids, has been determined to have 5 main gB genotypes: gB-1, gB-2, gB-3, gB-4, and gB-5. The most variable regions defining 5 genotypes are codons 26-70 which includes antigenic domain 2 (AD-2), and codons 441-490 that covers the furin cleavage site ^20–22^ (**Sup Fig. 1A, 1C**). A recent study analyzing the gB genotype variation between placebo recipients and vaccinees who received the gB/MF59 vaccine, which is based on the Towne strain (gB-1), showed that vaccinees were more likely to be infected with an HCMV strain encoding gB-3 and gB-5 compared to HCMV strains encoding gB-1, gB-2, and gB-4 genotypes via full gB open reading frame (ORF) next-generation sequencing (**Sup Fig. 1B**) ^23^. Yet, a previous report that examined the gB genotype of HCMV infections in this vaccine trial via Sanger sequencing did not reflect these findings ^24^. Thus, there is the potential that the gB/MF59 vaccine preferentially provided protection against certain HCMV gB genotypes, warranting examination if vaccine-induced immunogenicity can be broadened as a strategy to enhance vaccine efficacy.

Inclusion of multiple antigen valences and serotype-specific immunogens in a vaccine is commonly applied to improve vaccine immunogenicity via enhanced antigen-antibody interactions and to increase the vaccine breadth ^25^. Some examples of clinically-available multivalent vaccines include the poliovirus vaccine, pneumococcal vaccines, and human papillomavirus vaccine ^26^. In this study, we propose to include multiple gB genotypes in the nucleoside-modified mRNA-LNP vaccine platform to increase the vaccine protection breadth. While distinct serotypes recognized by polyclonal plasma of seropositive individuals have not been defined for HCMV, distinct gB genotype recognition has been described for gB-specific monoclonal antibodies ^27^. Thus, we hypothesized that a multivalent vaccine incorporating multiple gB genotypes could increase the anti-HCMV immunogenicity breadth and thus, a viable strategy for enhancing the efficacy of current HCMV vaccines.

Lipid nanoparticles-encapsulated nucleoside-modified mRNA (mRNA-LNP) is a novel vaccine platform that was translated for safe and effective human use in the severe acute respiratory syndrome coronavirus 2 (SARS-CoV-2) pandemic ^28^ and has had success in other vaccine pursuits, including human immunodeficiency virus type 1 (HIV-1)^29, 30^, Zika virus ^31^, influenza virus ^32^, and herpes simplex virus type 2 (HSV-2) ^33^. Additionally, the mRNA-LNP vaccine possesses multiple advantages, including a favorable safety profile, presentation of immunogens on the cell surface, rapid and cost-effective manufacturing process, and the ease of combining multiple antigens into a single vaccine ^34^. Indeed, the generation of multivalent mRNA-LNP vaccines proved to be a promising strategy for the development of broadly protective influenza virus vaccines ^35, 36^. Our lab previously compared the immunogenicity of the mRNA-LNP delivery platform to the protein-based full-length or ectodomain platform with squalene-like adjuvants encoding full-length HCMV gB genotype 1 (gB-1, sequence from Towne strain, GenBank accessioning number: FJ616285.1) in a rabbit model. Our data showed that the mRNA-LNP vaccine encoding full-length gB-1 enhanced the durability and breadth of peptide-binding responses ^37^. Recently, ModernaTX, Inc. has launched a phase III trial of an mRNA-LNP including HCMV Merlin strain gB and the pentameric glycoprotein complex after positive results in pre-clinical mice and non-human primate models (NCT05085366) demonstrated robust immunogenicity and epithelial cell neutralization responses ^38, 39^. The partial success of gB-based HCMV vaccines and the inclusion of gB in current HCMV vaccine candidates emphasize that gB remains a key vaccine target.

Here, we compare the immunogenicity of monovalent, bivalent, and pentavalent HCMV gB mRNA-LNP vaccines. Monovalent group 1 was administered gB1 mRNA-LNP. Bivalent group 2 was given gB1 + gB3 mRNA-LNPs, and pentavalent group 3 was vaccinated with gB1 + gB2 + gB3 + gB4 + gB5 mRNA-LNPs. New Zealand White rabbits were applied in this study due to their intermediate size and a longer life span, which allowed for large collection of blood samples and tissue biopsies from a single animal, investigation of the durability of vaccines ^40^, and has been validated to recapitulate Fc-mediated effector antibody responses mediated in primates ^41^. We evaluated whether bivalent and pentavalent vaccines including multiple gB genotypes could elicit broader protection against multi-genotypic HCMV infections than the monovalent HCMV gB immunogen. The findings in this study will be important for the guidance in improving the partial efficacy of HCMV gB-based vaccines.

## Results

### Study design and immunization schedule

The immunization schedule is shown in (**Fig. 1A**). A total of eighteen rabbits were divided into three groups (n=6 in each group) to receive gB mRNA-LNP monovalent, bivalent, and pentavalent vaccines, respectively, at weeks 0, 4, and 8. All three vaccines include a total of 50 µg (∼0.025 mg/kg) 1-methylpseudouridine-modified gB mRNA encoding single or multiple genotypes in each vaccine (Monovalent: 50 µg gB-1 mRNA; Bivalent: 25 µg gB-1+25 µg gB-3 mRNA; Pentavalent: 10 µg gB-1+10 µg gB-2+10 µg gB-3+10 µg gB-4+10 µg gB-5 mRNA.) These gB mRNA-LNP vaccines were given intradermally since this route showed the longest duration of mRNA translation and half-life compared to other routes^42^. To prevent the structural interference of the multivalent mRNA-LNP vaccines at the injection sites and to prevent administration differences among three groups, each rabbit received their vaccine doses split across 6 injection sites.

**Figure 1.**
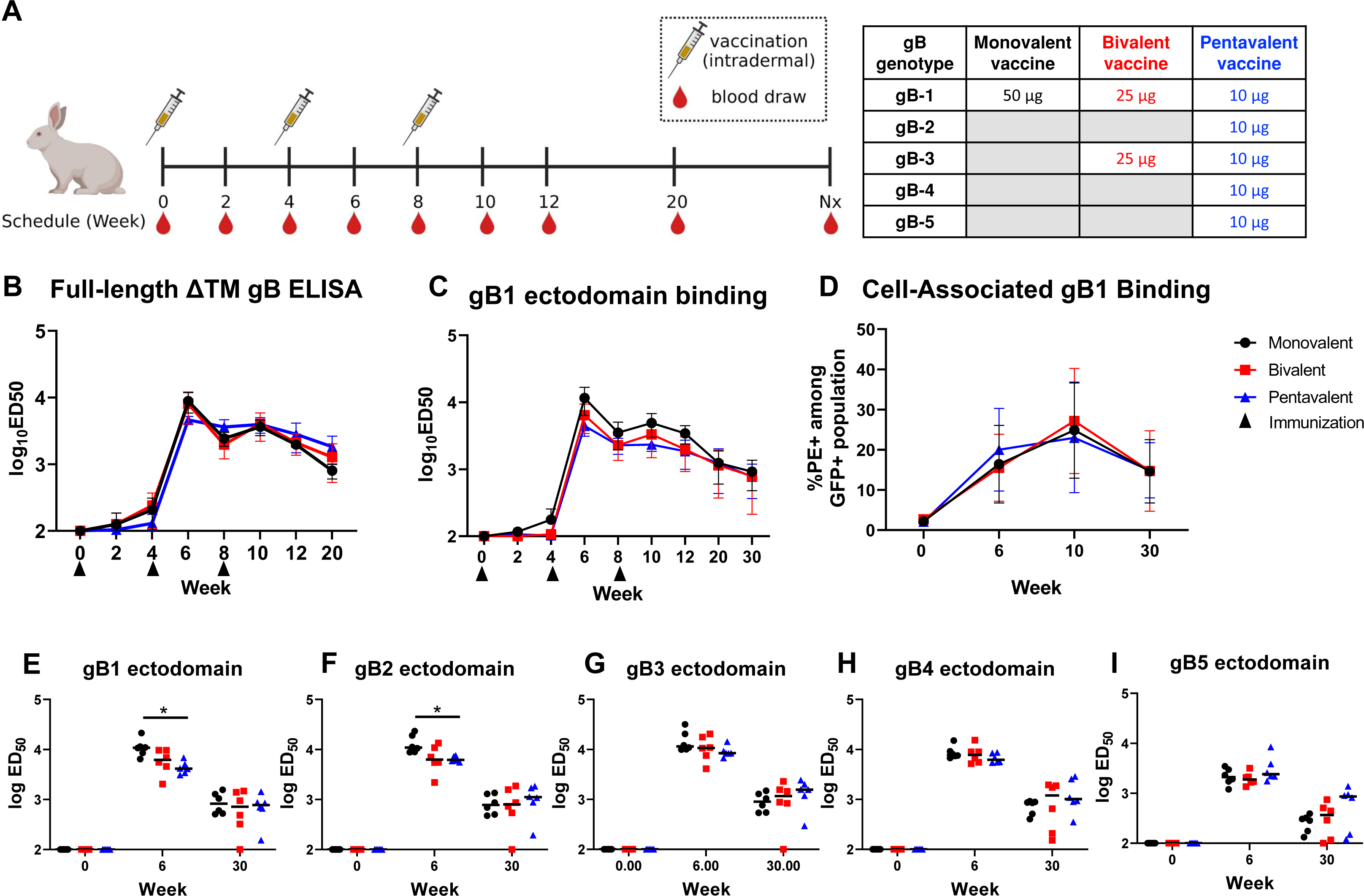
Monovalent, bivalent, and pentavalent gB mRNA-LNP vaccinated rabbit plasma IgG binding to soluble and cell-associated gB, and gB-specific IgG binding breadth. (A) Rabbit gB mRNA-LNP vaccination schedule. 18 rabbits were divided into three groups to receive gB mRNA monovalent, bivalent, and pentavalent vaccines, respectively, at week 0, 4, and 8. All three vaccines include a total of 50 µg 1-methylpseudouridine-modified gB mRNA while encoding single or multiple gB genotypes in each vaccine (Monovalent: 50 µg gB-1 mRNA; Bivalent: 25 µg gB-1+25 µg gB-3 mRNA; Pentavalent: 10 µg gB-1+10 µg gB-2+10 µg gB-3+10 µg gB-4+10 µg gB-5 mRNA.) Blood samples were collected every 2 weeks between week 0 and week 12, at week 20, and during necropsy. 15 animals underwent necropsy at week 30. The other 3 animals (1 animal from each group) received an extra boost at week 41 to prepare rabbit PBMCs for the functional antibody screening and underwent necropsy (Nx) at week 43. This figure was created with Biorender.com. (B-D) The dynamics of rabbit plasma IgG binding magnitude to soluble full-length gB with the deletion of transmembrane domain (B) and soluble gB-1 ectodomain (C) was measured by ELISA, and the IgG binding to cell-associated full-length gB-1 (D) for the pre-immune (week 0), peak immunogenicity (week 6 and 10), and durability (week 30) timepoints was estimated by gB-transfected cell binding assay. Data points are shown as the average IgG binding response with one standard deviation. A statistically significant difference was observed in the binding to the soluble gB-1 ectodomain (p-value 0.002, FDR adjusted p-value of 0.006). (E-I) Rabbit plasma IgG binding breadth to soluble gB ectodomains encoding 5 genotypes: gB-1 (E), gB-2 (F), gB-3 (G), gB-4 (H), and gB-5 (I) for the pre-immune (week 0), peak immunogenicity (week 6), and durability (week 30) timepoints was assessed by ELISA. Each data point represents the IgG binding response of one individual animal, with the median response labeled by a black line. Black circles: rabbits immunized with monovalent vaccine (gB-1); red squares: those immunized with bivalent vaccine (gB-1+gB-3); blue triangles: those immunized with pentavalent vaccine (gB-1+gB-2+gB-3+gB-4+gB-5). We first performed a multivariate Kruskal-Wallis (with p-values calculated via permutation). We observed a significant association at Week 6 and then followed up with Kruskal-Wallis tests at each gB. * indicate where there was a significant post-hoc association at week 6 (Holm adjusted p-value <0.10). Week 0 was not analyzed.

### Rabbit plasma gB-specific IgG binding to soluble gB protein, cell-associated gB, and linear gB peptide

We first measured the dynamics of rabbit plasma IgG binding to soluble full-length gB-1 without the transmembrane domain (ΔTM gB) and gB-1 ectodomain, matched to the monovalent vaccine strain (**Fig. 1B-C**). As expected, the IgG binding to both full-length gB-1 and gB-1 ectodomain has a peak at week 6 and 10, 2 weeks post the 2^nd^ and 3^rd^ boost, respectively. After the peak immunogenicity detected at week 6 and 10, the gB-specific IgG binding response waned gradually. Interestingly, although the bivalent and pentavalent vaccines include less gB-1 mRNA-LNP compared to the monovalent vaccine, no significant differences of the IgG binding magnitude to the soluble and cell-associated full-length gB-1 among three vaccine groups were observed (**Fig. 1B**). Notably, the monovalent vaccine elicited a statistically significantly different binding response to soluble gB-1 ectodomain in the longitudinal analysis, particularly at peak immunogenicity (higher at week 6 and 10), as expected. We next evaluated whether the bivalent and pentavalent vaccines could increase the breadth of IgG binding to multiple gB genotypes by comparing the rabbit plasma IgG binding to soluble gB ectodomains of 5 genotypes at week 0 (pre-vaccination), 6 (peak immunogenicity), and 30 (durability timepoint) (**Fig.1E-I)**. The IgG binding to gB-1, gB-2, gB-3, and gB-4 ectodomains was approximately 10-fold higher than that to gB-5 ectodomain at week 6 among all groups, while the IgG binding magnitude at week 30 was similar against 5 genotypes. The monovalent vaccine group elicited a statistically higher IgG response against gB-1 and gB-2 compared to the other two groups at week 6. For the durability timepoint at week 30, although the pentavalent vaccine elicited a slightly higher IgG binding to gB-5 ectodomain in some animals compared to the monovalent and bivalent vaccine, no significant differences among three groups emerged.

A previous study from our lab demonstrated that the gB/MF59 vaccine protection against HCMV acquisition was associated with the level of plasma IgG binding to cell-associated gB. In this study, it was also shown that the conformation of gB expressed in soluble form is different from that expressed on cell surface ^6^. Thus, we also assessed the dynamics of rabbit plasma IgG binding to cell-associated gB-1 (**Fig. 1D**) and the other gB genotypes **(Sup Fig.2C-E)**. To measure the IgG binding to cell-associated gB of different genotypes, we designed green fluorescence (GFP)-tagged plasmids that include full-length gB sequences encoding five genotypes and T2A peptide sequence to prevent the structural interference between gB and GFP (gB-T2A-GFP plasmid). Unlike the IgG binding to soluble gB, the IgG binding to cell-associated gB-1 peaked at week 10 of the vaccine schedule, 2 weeks post the 3^rd^ boost. No significant differences of the plasma IgG binding kinetics to cell-associated gB-1 were observed among three groups **(Fig.1D)**. Due to the varied transfection efficiency, the IgG binding against five gB genotypes at week 6, 10, and 30 was normalized to that at week 0 **(Sup Fig.2C-E)**. No significant differences of IgG binding to cell-associated gB encoding five genotypes were observed among three vaccine groups.

Further, we measured the rabbit plasma IgG binding response to linear, overlapping 15-mer gB peptides covering the variable region, amino acid codons 1-77 of gB-1 or gB-2/3 genotypes (27 unique peptides) (**Sup Table 1**). We designed peptides based on the monovalent vaccine-matched genotype, gB-1, and the genotypes only included in the bivalent and pentavalent vaccine, gB-2/3, since gB-2 and gB-3 sequences share the greatest similarity compared to other genotypes in this region (**Sup Fig. 1A**). The gB peptide-binding IgG response of the monovalent, bivalent, and pentavalent vaccinated rabbit plasma at week 10 revealed no observable differences comparing the number of peptides bound by plasma IgG from gB-1 or gB-2/3 among three vaccine groups (**Sup Fig. 3B-C**). The bivalent and pentavalent gB mRNA LNP vaccines had similar IgG binding breadth compared to the monovalent mRNA LNP vaccine.

### Neutralization and antibody-dependent cellular phagocytosis (ADCP) responses

With the incorporation of multiple gB genotypes in the multivalent vaccines, we hypothesized that bivalent and pentavalent vaccines could elicit a broader functional antibody response including neutralizing antibody and antibody-dependent phagocytosis (ADCP) response against HCMV strains encoding different gB genotypes. We first investigated the vaccine-elicited neutralizing antibody response against the Towne, AD169r, and Toledo strains encoding gB-1, gB-2, and gB-3, respectively, in fibroblasts **(Fig. 2A-C, 2E-G)** and epithelial cells **(Fig. 2D, 2H)**. A previous study showed that complement enhances and can be required for the neutralizing potency of gB-specific antibodies in a rabbit model. Therefore, the vaccine-elicited neutralizing antibody response was assessed with and without rabbit complement here ^43^. As expected, the neutralizing titer without rabbit complement was lower and mostly undetectable against AD169r and Toledo strain compared to that performed in the presence of rabbit complement **(Fig. 2E-H)**. Therefore, we did not perform the statistical analysis on the neutralizing antibody titer data generated without rabbit complement. We utilized a Kruskal-Wallis test (p-values determined via 100,000 permutations) to determine whether the neutralizing antibody response with rabbit complement among three vaccine groups is different. Inconsistent with our expectation, at peak immunogenicity (week 6 or 10), the monovalent vaccine group exhibited a higher or similar neutralization capability against Towne, AD169r, and Toledo strain compared to the multivalent vaccine groups by the multiple testing correction analysis. After performing the post-hoc analysis (15 Wilcoxon Rank Sum tests), we observed that the monovalent vaccine elicited a statistically higher neutralizing response against Toledo strain at week 10 than the bivalent vaccine and against the Towne at week 10 compared to the pentavalent vaccine. Additionally, no statistical differences of the neutralizing antibody responses in epithelial cells against the AD169r strain, which was repaired in the ULb’ region to maintain epithelial cell tropism, were observed. These results suggest that the bivalent and pentavalent vaccines did not elicit a broader neutralizing antibody response at peak immunogenicity as we expected. Interestingly, the bivalent and pentavalent vaccine elicited a statistically higher neutralizing titer than the monovalent vaccine at durability time point (week 30) against Toledo strain (a gB-3 HCMV strain included in the bivalent and pentavalent vaccine design) in the presence of complement in fibroblasts. Due to the sample size and number of post-hoc tests, However, no statistical significance was observed in post-hoc analysis comparing monovalent and bivalent (p-value 0.065, p_Holm_ 0.39), or monovalent and pentavalent (p-value 0.009, p_Holm_ 0.11) vaccine groups. We also performed a secondary analysis by Wilcoxon rank sum test to compare the monovalent vaccine to the multivalent vaccines (combine bivalent and pentavalent vaccine groups) and observed a statistically higher neutralizing titer against AD169r strain in fibroblasts at week 30 (not indicated in the figure).

**Figure 2.**
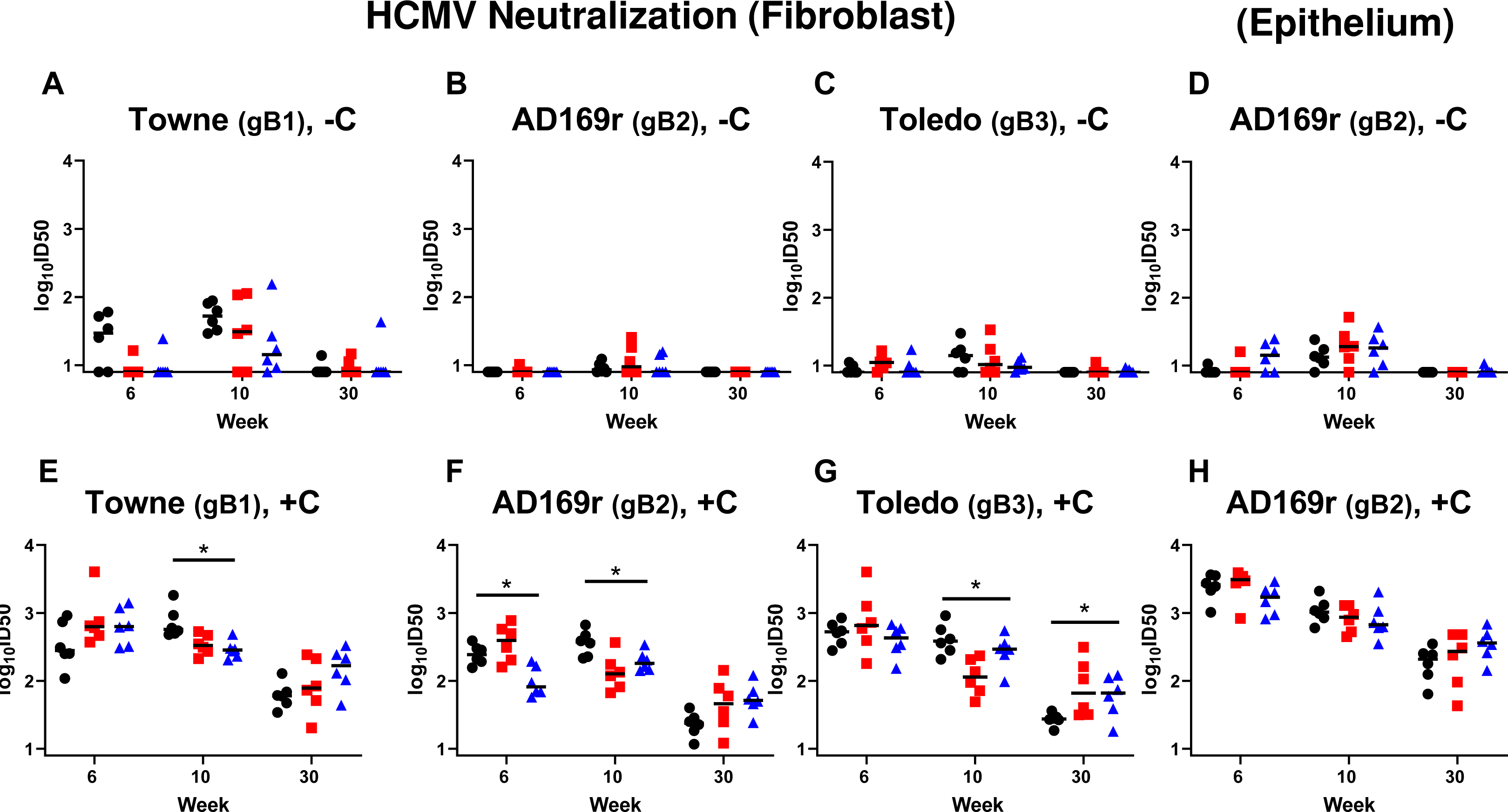
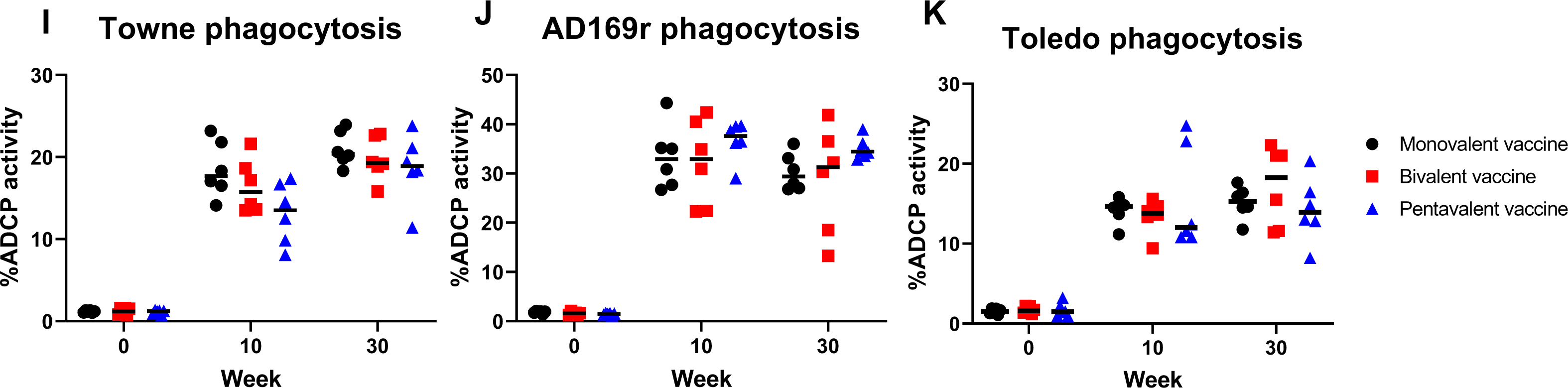
Monovalent, bivalent, and pentavalent gB mRNA-LNP vaccine elicit similar magnitude and breadth of HCMV-neutralizing antibody response against HCMV strains with gB genotype 1, 2, or 3. Rabbit plasma IgG neutralization of HCMV Towne strain encoding gB-1 (A,E), AD169r strain encoding gB-2 (B,D, F, H), and Toledo strain encoding gB-3 (C, G) without the addition of purified rabbit complement (-C) (A-D) or with the purified rabbit complement (+C) (E-H) was estimated in fibroblasts (A-C, E-G) or epithelial cells (D,H).). We did not test the without complement data (-C, Figure 2A-2D). *, P<0.1, Kruskal-Wallis test performed with the multiple testing correction via FDR. Post-Hoc tests were done via Wilcoxon Rank Sum Test to assess for differences between pairs of vaccine groups. Phagocytosis of the whole HCMV virion of the Towne strain (I), AD169r strain (J), and Toledo strain (K) was measured by flow cytometry. Antibody neutralization and phagocytosis activity was assessed at the pre-immunization (week 0), peak immunogenicity (week 6, 10), and durability (week 30) timepoints. Each data point represents the neutralization activity or whole virion phagocytosis response of one individual animal, with the median response labeled by a black line. Black circles: rabbits immunized with monovalent vaccine; red squares: those immunized with bivalent vaccine; blue triangles: those immunized with pentavalent vaccine. *, P<0.1, Kruskal-Wallis test performed with multiple testing correction by FDR. Post-Hoc comparisons were done as for Figures 2E - 2H.

Subsequently, we examined whether a multivalent gB mRNA vaccine could increase the breadth of non-neutralizing antibody functions. Our lab previously showed that gB-specific antibodies elicited by the gB/MF59 vaccine mediated robust ADCP response against whole HCMV virions at the similar level to those elicited by chronically HCMV-infected individuals^5^. In addition, high magnitude ADCP response against HCMV virions was shown to be associated with a lower risk of congenital HCMV transmission in HCMV seropositive women ^44^. Therefore, we hypothesized that the inclusion of multiple gB genotypes in the vaccine design could increase the antibody breadth and subsequently enhance the ADCP response. We measured the rabbit plasma ADCP response against the whole HCMV virions from the Towne, AD169r, and Toledo strains encoding gB-1, gB-2, and gB-3, respectively (**Fig. 4I-K**). The anti-HCMV ADCP potency of vaccinated-rabbit plasma IgG peaked at week 10 and showed a durable response until week 30. However, the bivalent and pentavalent vaccines did not elicit a higher ADCP response against HCMV virions encoding gB-1, gB-2, or gB-3 as we expected. Therefore, our data suggest the multivalent vaccines might not show an advantage for the breadth of this key Fc-mediated effector function.

### Monoclonal antibody isolation from rabbit PBMCs

After characterizing mono-and multivalent vaccine-elicited plasma IgG binding and functional antibody responses, we further investigated whether rabbits immunized with the multivalent vaccines could generate gB genotype-specific monoclonal antibodies (mAbs). We performed single memory B cell culture from rabbit peripheral blood mononuclear cells (PBMCs) at week 10 (2 weeks post 3^rd^ boost, peak immunogenicity) to isolate gB-specific mAbs, using methods previously reported^45^. Due to the limited cell numbers, we mixed two rabbit PBMCs samples from the monovalent and the pentavalent group to perform single memory B cell culture. From the 600 single memory B cell cultures, we identified 38 mAbs in culture supernatant that bound to full-length gB-1 **(Sup Fig. 5A-B)** and 1 of those 38 (mAb 2M7) potently neutralized HCMV in fibroblasts against AD169r strain. The potent neutralizing mAb 2M7 was further cloned and purified for the downstream antibody characterization **(Fig. 3A)**.

**Figure 3.**
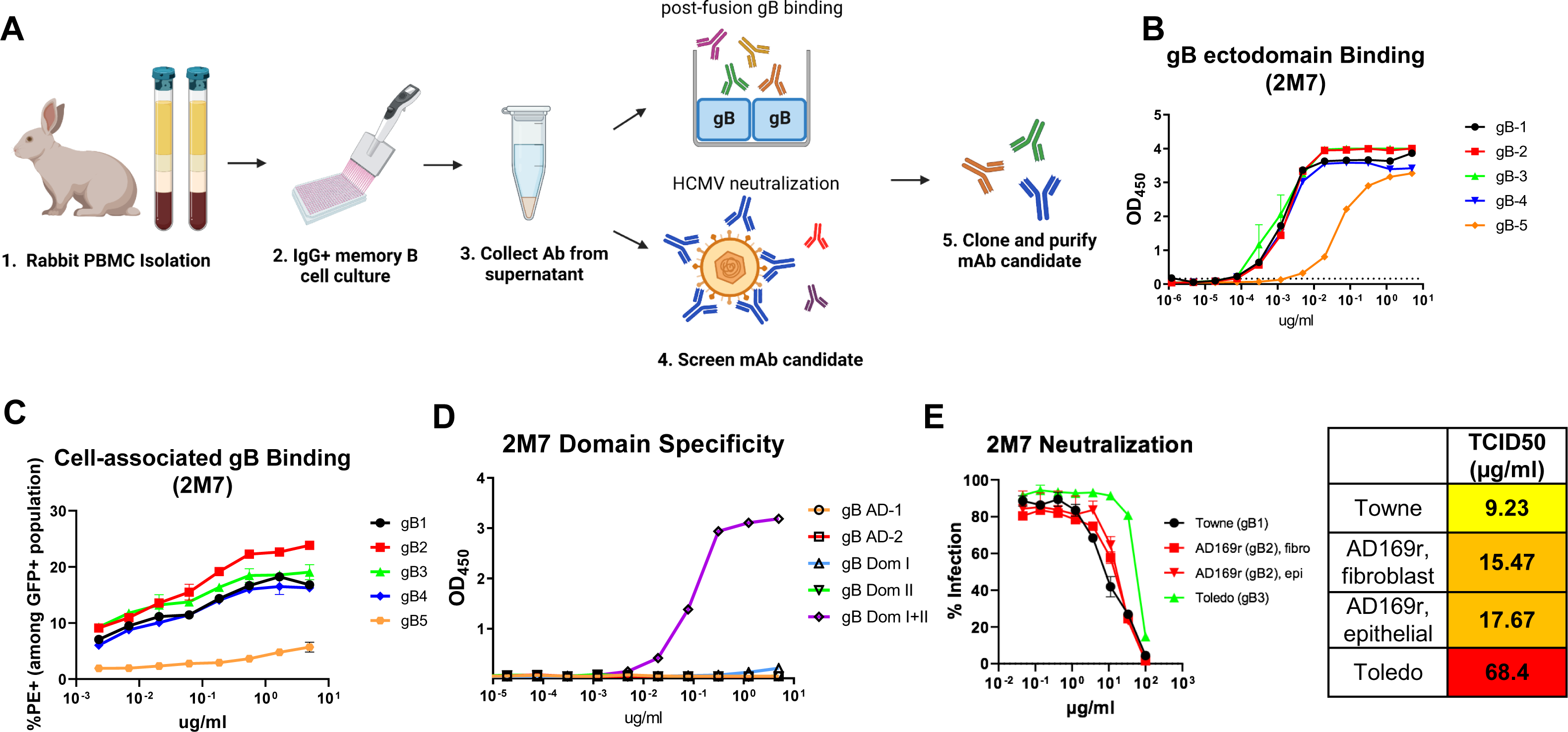
Genotype-dependent binding and neutralization properties of a gB-specific monoclonal antibody isolated from transformed memory B cells of multivalent gB mRNA-LNP-immunized rabbits. (A) gB-specific mAb was isolated from rabbit PBMCs memory B cell culture. Rabbit PBMCs at peak immunogenicity (week 10) were isolated from EDTA whole blood and the IgG+ memory B cells were cultured. The memory B cell culture supernatant was collected and performed post-fusion gB binding ELISA or fibroblast neutralization against AD169r strain for screening. The mAb candidate with potent gB-binding was cloned and purified for the downstream characterization. (B-C) Isolated gB-specific mAb 2M7 binding to soluble gB ectodomains encoding 5 genotypes was measured by ELISA (B) and cell-associated full-length gB encoding 5 genotypes was estimated by flow cytometry (C). 2M7 binding to gB genotypes is color-coded in lines: black, gB-1; red, gB-2; green, gB-3; blue, gB-4; orange, gB-5. (D) 2M7 binding strength to gB AD-1, AD-2 site 1, Domain I, Domain II, and Domain I+II was measured by ELISA, demonstrating conformational-dependent Dom I+II specificity. 2M7 binding to gB domains is color-coded with symbols: black circle, AD-1; red square, AD-2 site 1; green upward triangle, Domain I; blue downward triangle, Domain II; orange diamond, Domain I+II. (E) The neutralization capability of 2M7 against HCMV Towne strain (gB-1) and Toledo strain (gB-3) was estimated in fibroblasts and that against AD169r strain (gB-2) was measured in fibroblasts and epithelial cells. 2M7 neutralization is color-coded based on the HCMV strains: black, Towne strain; red, AD169r strain; green, Toledo strain.

We characterized the genotype-dependent binding and neutralization property of mAb 2M7 against HCMV strains encoding different gB genotypes **(Fig. 3B-E)**. The mAb 2M7 demonstrated a similar binding to gB ectodomain or cell-associated gB encoding genotypes 1-4, but a lower binding to gB-5 **(Fig. 3B-C)**. We further characterized the epitope binding specificity of 2M7 and found that 2M7 demonstrated a high binding to soluble gB domain I+II construct but not to AD-1, AD-2, domain I, and domain II, individually **(Fig. 3D).** Further, 2M7 exhibited the highest neutralizing potency against the HCMV Toledo strain (gB-3) when compared to the Towne and AD169r strains (gB-1 and gB-2, respectively, **Fig. 3E)**. These results demonstrate that the mAb 2M7 possess a gB genotype-dependent function.

### Vaccine-elicited gB-2-specific PBMC and splenic T cell response

T cell immunity is crucial for controlling HCMV infection as well as generating broad antibody response, which is commonly assessed as a part of the immunogenicity of HCMV vaccine candidates ^46, 47^. Thus, a broad T cell response across gB genotype strains would also be desirable in HCMV vaccine candidates. We hypothesized that the pentavalent vaccines could elicit a more robust T cell response against a gB genotype not included in the monovalent or bivalent vaccine, for instance, gB2. To test our hypothesis, we stimulated rabbit PBMCs at peak immunogenicity (week 6) and splenocytes at necropsy (week 30) with HCMV gB-2 peptide pools and measured the IFN-γ-secreting cell numbers by ELISPOT (**Fig. 4A-B, Sup Fig. 6A-B**). The pentavalent-vaccinated rabbit PBMCs demonstrated the highest IFN-γ T cell response against gB-2 at week 6, followed by those from the bivalent and monovalent vaccine groups (**Fig. 4A**). This result is consistent with our expectation that the pentavalent vaccine is the only vaccine that includes gB-2 immunogen among three groups. By performing the post-hoc analysis, a statistically higher IFN-γ-secreting T cell response was observed comparing the bivalent and monovalent, the pentavalent and monovalent vaccine groups, respectively. No statistical significance of IFN-γ-secreting splenic T cell response against gB-2 was observed among three vaccine groups. Interestingly, the bivalent vaccine elicited a higher IFN-γ splenic T cell response against gB2 at week 6 and 30 than the monovalent vaccine group (**Fig. 4B**), despite that gB2 was not a component of either vaccine.

**Figure 4.**
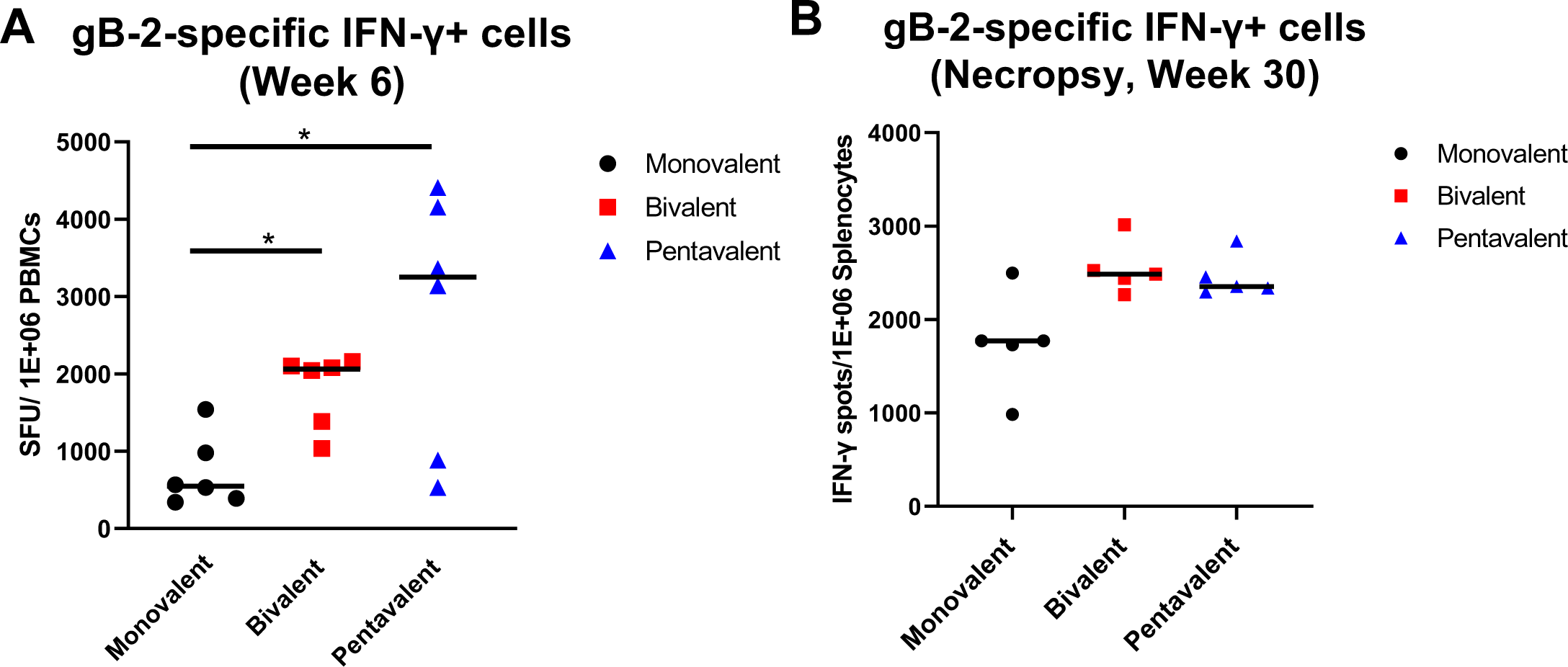
Multivalent gB mRNA-LNP vaccine elicited a stronger gB-2-specific (AD169r strain) IFN-r+ cell response in PBMCs at peak immunogenicity and in spleen at necropsy than monovalent gB mRNA-LNP vaccine. gB-2-specific IFN-r+ cells from rabbit PBMCs or splenocytes were measured by ELISPOT. Full length, overlapping gB-2 peptide pool stimulation was performed in duplicate. (A) Quantification of IFN-r+ spots from rabbit PBMCs stimulated with gB-2 peptides at peak immunogenicity (Week 6) from monovalent, bivalent, and pentavalent vaccine group. (B) Quantification of IFN-r+ spots from rabbit splenocytes stimulated with gB-2 peptides at necropsy from monovalent, bivalent, and pentavalent vaccine group. 15 animals underwent necropsy at week 30, while 3 animals (1 animal from each group) underwent necropsy at week 43 due to an extra boost prior to B cell isolation at week 41. Each data point represents one peptide-stimulated well from individual animals, with the lines designating the median. Black circles: rabbits immunized with monovalent vaccine; red squares: those immunized with bivalent vaccine; blue triangles: those immunized with pentavalent vaccine. . **, P<0.01, ***, P<0.001, Kruskal-Wallis test with Bonferroni correction. An association was observed at week 6 (unadjusted p-value of 0.025). Post-Hoc tests via Wilcoxon Rank Sum revealed it was largely driven by differences between Monovalent and the two other vaccines groups.

## Discussions

Our study is the first to investigate whether the inclusion of multiple gB genotypes in HCMV vaccine design would provide broader immunity and lead to better protection against multiple HCMV strains. Applying the recently translated and highly successful nucleoside-modified mRNA-LNP platform, we vaccinated three groups of rabbits with the monovalent (gB-1), bivalent (gB-1, gB-3), and pentavalent (gB-1, gB-2, gB-3, gB-4, and gB-5) gB vaccines, respectively, and compared the humoral and cellular immunity among three groups. The kinetics of rabbit plasma IgG binding response against full-length soluble and cell-associated gB-1 post immunization was similar among three groups. Yet, the multivalent vaccine groups did not demonstrate broader IgG response against soluble and cell-associated gB encoding five genotypes or enhance the breadth of functional antibody responses. Interestingly, the multivalent vaccines elicited a higher T cell magnitude than monovalent vaccine against the monovalent and bivalent vaccine-mismatched gB genotype (gB-2) at peak immunogenicity. This result implies that the multivalent gB mRNA vaccine might have an advantage for increasing T cell breadth but requires further validation.

The concept of multivalent antigen presentation has been practiced as a common strategy to increase the vaccine protection breadth. Several multivalent vaccines have been licensed in the U.S., including pneumococcal vaccines and human papillomavirus (HPV) vaccines ^25, 26, 48^. Two pneumococcal multivalent vaccines targeting high risk strains of the approximately 100 identified have been licensed, pneumococcal polysaccharide vaccine (PPSV23) and pneumococcal conjugate vaccine (PCV13) ^49, 50^. As the major cause of cervical cancer, HPVs have more than 200 distinct types that could be divided into high-risk or low-risk groups. Currently, three multivalent HPV vaccines targeting high-risk HPV types have been licensed in the U.S, with the 9-valent vaccine being the most highly distributed.

We hypothesized that the multivalent gB mRNA-LNP vaccine could increase the HCMV vaccine protection breadth and thus the efficacy based on the success of multivalent pneumococcal and HPV vaccines. Our previous work demonstrated that the nucleotide-modified mRNA-LNP gB vaccine elicited a more durable and broader antibody response compared to the gB protein subunit vaccine^51^. Additionally, in an influenza-specific antibody characterization study, it was shown that the antibody breadth necessitated the interactions between Fc domain and Fcγ receptor ^52, 53^. This interaction is the initiation step for Fc-mediated effector responses that was shown to be critical for protection against HCMV, such as ADCP response. We designed an HCMV vaccine to include the full-length gB mRNA rather than only include the variable regions to maximize the immunogenicity since there were five antigenic domains (AD) identified across the HCMV gB sequence, AD-1 to AD-5 ^54–58^. We chose to test the multivalent gB mRNA vaccine efficacy in a rabbit model as we previously validated that the *in vitro* assays could be applied to measure the vaccine-elicited functional antibody responses and epitope specificity ^41^. We also chose to immunize rabbits intradermally instead of intramuscularly to achieve the maximum vaccine immunogenicity. While the clinical SARS-CoV-2 mRNA-LNP vaccines were delivered intramuscularly since this route is easier to implement, a previous mRNA-LNP vaccine route of administration study using a mouse model found the intradermal route showed the longest duration and half-life of mRNA translation compared to other routes^42^, providing the most sensitive assessment of vaccine immunogenicity. The intradermal route has also been applied in the rabbit models in multiple mRNA-LNP vaccine studies ^29, 32, 51^.

Our data showed that the pentavalent vaccine that included only a fifth of the total gB-1 mRNA-LNP dose compared to that of the monovalent vaccine elicited a similar binding and functional antibody response than the monovalent vaccine. The multivalent vaccines did not increase the breadth of IgG binding breadth or functional neutralizing and non-neutralizing antibody responses as we expected. These findings suggest that the gB mRNA-LNP vaccine-elicited humoral response mainly targets the conserved domains instead of variable regions on the gB sequence. Additionally, it may be that the structural differences among five gB genotypes are subtle, so the multivalent vaccine did not elicit a polyclonal response that had distinct breadth from the single valent vaccine. This conclusion could be supported by ∼96% mean conserved identity at amino acid level among gB genotypes^59^. However, this conclusion is opposite to the inference drawn by the identification of genotype-specific mAbs isolated from the natural HCMV-infected healthy adults in a previous study^27^. Currently, only a gB-2 (AD169r) ectodomain post-fusion structure and gB-1 (Towne) pre-fusion structure have been published^60, 61^. Although studies regarding multiple gB genotyping have been widely published ^62^, the structural differences among gB genotypes, particularly the flexible AD2 domain, remain elusive.

Interestingly, while multivalent gB vaccination did not elicit a polyclonal response that had distinct breadth from single valent vaccination, the recognition of distinct gB genotypes was demonstrated on the mAb level. Previously, we have compared the gB-specific antibody binding preference to five genotypes from the HCMV-infected individuals ^27^. In this study, we isolated a gB-specific neutralizing antibody 2M7 from vaccinated rabbit PBMCs that demonstrated genotype-dependent function. The mAb 2M7 showed a similar binding to soluble and cell-associated proteins encoding gB-1, gB-2, gB-3, and gB-4 genotypes, but a lower binding to gB-5 soluble and cell-associated proteins. This finding suggests the gB-5 protein structure might be the most dissimilar among five genotypes, even though the gB-3 and gB-5 share a greater genetic similarity in the amino acid sequence ^23, 24^. Additionally, 2M7 neutralized the HCMV Toledo strain (gB-3) the most potently although also neutralized HCMV Towne (gB-1) and AD169r (gB-2) strain. The binding and neutralization profile suggests that 2M7 might target variable regions of gB. Therefore, we characterized the 2M7 binding domain specificity. Interestingly, we found that 2M7 exhibited the strongest binding to gB Dom I+II but no binding to individual AD-1, AD-2, Dom I, and Dom II. Concordant with a previous study, gB Dom I and Dom II were found to be the major targets of gB-specific neutralizing antibodies ^58^. Further characterization of gB-specific genotype-dependent neutralizing antibodies might direct the future HCMV gB vaccine design.

Our data also showed that a multivalent gB vaccines elicited a higher gB-2-specific T cell response compared to that of monovalent vaccine, which did not contain the gB2 antigen. As expected, the pentavalent vaccine elicited the highest IFN-γ-secreting T cell response among three vaccine groups at peak immunogenicity. Yet, surprisingly, the bivalent vaccine, which did not contain the gB2 antigen, also demonstrated a higher T cell response than the monovalent vaccine at peak immunogenicity and durability timepoints. Considering the inclusion of gB genotypes in the monovalent (gB-1) and bivalent (gB-1 and gB-3) vaccines, this observation might indicate the T cell epitopes of gB-2 and gB-3 genotypes are somewhat similar compared to gB-1. The sequence alignment suggests that gB-2 and gB-3 are almost identical at codon 28-70 while gB-1 has completely distinct codons at the same region. Therefore, the codon 28-70 region potentially contains a T cell epitope target, although a previous study showed the gB-specific T cell response elicited by HCMV-seropositive healthy donors did not target peptides from the first 80 amino acids ^63^. The higher bivalent vaccine-elicited T cell response also suggests the vaccine-elicited T cell response breadth may increase even if only including two genotypes in HCMV vaccine design. As the T cell response breadth was not the main outcome of this study that sought to generate breadth in anti-HCMV functional antibody responses, further studies are required to identify whether including multivalent immunogens could increase the breadth of vaccine-elicited T cell responses against HCMV infection.

Limitations of this study include the animal model and the species specificity of HCMV that complicates challenge studies. Compared to rodents, the rabbit model has multiple advantages, including their intermediate size for large sample collection, long life span for vaccine boosts, lack of an inbred strain to better mimic human populations, and a closer phylogenetic relationship with human beings^40, 64–68^. Although rabbits are commonly applied in preclinical vaccine studies, the immune system of rabbits may not represent human B and T cell repertoire. We did not examine whether the multivalent vaccines expand the T cell repertoire, partly due to the cross-species model limitations, and therefore this should be studied in future human studies. Further, the small animal numbers in each group make it less possible to detect the subtle differences from the vaccine-elicited polyclonal immunity. Finally, a challenge model cannot be applied in the setting of HCMV vaccine preclinical testing due to the species-specificity of the HCMV variants. Thus, we chose to compare the breadth of the responses between monovalent and multivalent vaccines against HCMV strains as the primary outcome of this study.

Future HCMV vaccine design should consider several perspectives. We recently demonstrated that elite neutralizers of HCMV have potent neutralizing antibody responses that target multiple glycoprotein specificities ^69^, including pentameric and trimeric glycoprotein complexes on the HCMV virion surface. However, there was no difference in gB-specific binding breadth comparing the HCMV-seropositive elite neutralizers to non-elite neutralizing individuals. Further, the neutralizing antibody response may not be the only antibody function that determines the HCMV vaccine efficacy. Since non-neutralizing antibody responses were shown to be associated with a decreased risk of congenital HCMV transmission and the vaccine efficacy, future HCMV vaccines should target key non-neutralizing antibody functions such as ADCP and antibody-dependent cellular cytotoxicity (ADCC) ^5, 70^. In fact, a recent paper showed that HCMV non-structural glycoproteins are the major ADCC targets^71^. This finding suggests the future HCMV vaccine design that aims to increase the polyfunctional antibody response breadth should focus on the inclusion of additional glycoproteins beyond gB, as well as non-neutralizing antibody targets, in order to enhance the vaccine efficacy.

Overall, our study demonstrates that the multivalent gB mRNA-LNP vaccine did not increase the breadth of the gB-specific humoral response. This question of multi-vs single-valent anti-viral vaccines has become highly relevant in the design and roll out of the next generation SARS-CoV-2 mRNA-LNP vaccines to include the Omicron variant. A recent study showed that the SARS-CoV-2 mRNA-LNP vaccine boost in naïve individuals elicited an increased plasma neutralizing antibody response but a variable antibody response breadth compared to the convalescent individuals ^72^. Since the inclusion of the Omicron variant provided minimal gains in protective efficacy against new SARS-CoV-2 strains, their impact on the SARS-CoV-2 pandemic remains unknown ^73–76^. While the B cell response breadth against HCMV gB did not benefit from multivalent vaccines in these studies, a higher magnitude T cell response was elicited against monovalent vaccine-mismatched gB genotype, implying that the multivalent HCMV glycoprotein immunization should be considered as a strategy for those vaccines that seek to enhance T cell response breadth. With the ease of including multiple antigens in future mRNA-LNP vaccines, studies should continue to assess the ideal diversity and quantity of antigens that can be administered concurrently in a vaccine to achieve the most effective immunity for CMV as well as other pathogens.

## Materials and Methods

### gB mRNA production and formulation in lipid nanoparticles

The modified mRNAs encoding HCMV gBs were produced as previously described using T7 RNA polymerase (MEGAscript; Ambion) on codon-optimized linearized plasmids ^77, 78^. The gB sequences are below: gB-1 from Towne strain (GenBank accession number: ACM48044.1; gB-2 from AD169 strain, GenBank accession number: DAA00160.1), gB-3 (Toledo strain, GenBank accession number: ADD39116.1), gB-4 (C194 strain, GenBank accession number: AAA45925), and gB-5 (saliva isolate, GenBank accession number: AZB53144). The mRNAs were transcribed to contain 101-nucleotide-long poly(A) tails. To generate modified nucleoside-containing mRNA, m1Ψ-5’-triphosphate (TriLink) was used instead of UTP. The mRNAs were then capped using an m7G capping kit with 2’-O-methyltransferase (CellScript). Th gB-1 mRNA was purified by fast protein liquid chromatography (FPLC; Akta purifier; GE Healthcare)^79^, the gB-2, gB-3, gB-4 and gB-5 mRNAs were purified by cellulose-based purification ^80^. All mRNAs were analyzed by electrophoresis using agarose gels, and stored at -20°C. The purified m1Ψ-containing HCMV gB mRNAs were encapsulated in LNPs using a self-assembly process. This process involved a rapid mixture of mRNA in pH 4.0 aqueous solution and lipids dissolved in ethanol solution. The LNPs used in this study were similar in composition to those described previously ^81, 82^. These LNPs contain an ionizable cationic lipid (proprietary to Acuitas), phosphatidylcholine, cholesterol, and polyethylene glycol-lipid. The proprietary lipid and LNP composition are described in US patent US10,221,127. They had a diameter of ∼80 nm, as measured by dynamic light scattering using a Zetasizer Nano ZS (Malvern Instruments Ltd.) instrument.

### Animal handling, immunization, sample collection and processing

Adult New Zealand White rabbits (9-10 months old) were a kind gift from Dr. Herman Staats (Duke University) and housed at Duke University. Rabbits were previously utilized to evaluate the nasal immunogenicity of an unrelated antigen. The immunization of the gB mRNA-LNP vaccines was via intradermal injections on the neck. The monovalent vaccine group was given a total of 6 injections with 8.3 µg gB-1 mRNA-LNP; the bivalent vaccine group was given 3 injections with 8.3 µg gB-1 mRNA-LNP and 8.3 µg gB-3 mRNA-LNP; the pentavalent vaccine group was given 1 injection with 10 µg gB-1, 10 µg gB-2, 10 µg gB-3, 10 µg gB-4, 10 µg gB-5, and phosphate buffered saline (PBS), respectively. For blood collections from central ear artery, animals were first sedated with 1mg/kg of body weight acepromazine subcutaneously and the ears were applied with 1% lidocaine topically. The blood was collected via auricular venipuncture and stored in an EDTA-anticoagulated tube before samples processing. Rabbit plasma was separated from whole blood by centrifugation and stored in -80C prior to running assays. Peripheral blood mononuclear cells (PBMCs) were isolated by density gradient centrifugation using Lymphocyte-Mammal cell separation medium (Cedarlane Laboratories).

For necropsy, the animals were euthanized using 0.5 ml of subcutaneously injected xylazine (100 mg/ml) plus ketamine (500 mg/ml), mixed together in a 1:5 ratio, followed by 0.5 ml of intracardiac pentobarbital sodium plus phenytoin sodium (Euthasol). Blood, spleen, and bone marrow samples were collected at necropsy. The collection of 50ml rabbit blood was obtained from cardiac puncture with a 21-gauge needle. Splenocytes were isolated from the spleen by manual tissue disruption and crushing through a 100 µm cell strainer, followed by density gradient centrifugation using Lymphocyte-Rabbit cell separation medium (Cedarlane Laboratories). Bone marrow samples were collected from femurs and flushed through a 100 µm cell strainer.

### Cell Culture

Human epithelial kidney 293T cells (ATCC) were maintained in Dulbecco’s modified Eagle medium (DMEM) supplemented with 10% fetal bovine serum (FBS), 25mM HEPES, and Pen-Strep. Human retinal pigment epithelial (ARPE-19) cells (ATCC) were maintained in DMEM supplemented with 10% FBS and Pen-Strep. Human foreskin fibroblast HFF-1 cells (ATCC) were maintained in DMEM supplemented with 20% FBS, 2mM L-glutamine, 25mM HEPES, Pen-Strep, and 50 µg/ml Gentamicin. Human monocytes THP-1 cells (ATCC) were maintained in Roswell Park Memorial Institute Medium (RPMI) 1640 media supplemented with 10% fetal bovine serum (FBS). Expi293F™ cells (Thermo Fisher Scientific) were cultured in Expi293F™ expression medium (Thermo Fisher Scientific).

### Protein Production

The histidine (His)-tagged soluble gB ectodomain plasmids were designed by removing the transmembrane and cytosolic domains of gB sequences for five genotypes, respectively, and cloned into pcDNA3.1(+) mammalian expression vector (Invitrogen). The gB sequences of the five genotypes are below: gB-1 from Towne strain (GenBank accession number: ACM48044.1; gB-2 from AD169 strain, GenBank accession number: DAA00160.1), gB-3 (Toledo strain, GenBank accession number: ADD39116.1), gB-4 (C194 strain, GenBank accession number: AAA45925), and gB-5 (saliva isolate, GenBank accession number: AZB53144). Expi293F™ cells were transiently transfected with His-tagged gB ectodomain plasmids for five days and later harvested with Nickel-NTA resin (Thermo Fisher Scientific).

The protein production and purification of gB Domain I, Domain II, and Domain I+II was previously reported^5^. In brief, sequences encoding HCMV Merlin strain gB Domain I (codon 133-343), Domain II (codon 112-133, 343-438) and gB Domain I+II (codon 112-438) were tagged with hemagglutinin (HA) tag at the 5’ end, Domain I and Domain I+II were also tagged with an avidin and poly-his tag at the 3’ end. Since gB Domain II was discontinuous, the two segments within gB Domain II were joined by the flexible linker. These gB domain sequences were cloned into pcDNA3.1(+) mammalian expression vector (Invitrogen) and transiently transfected into Expi293F™ cells for five days. The gB Domain I and Domain I+II were purified using Nickel-NTA resin (Thermo Fisher Scientific), and the gB Domain II was purified with lectin resin (VWR.)

### Rabbit plasma IgG binding to soluble gB proteins and mAb binding specificity to gB domains by ELISA

To estimate the rabbit plasma IgG binding to soluble gB proteins, 384-well clear-bottom ELISA plates (Corning) were coated overnight at 4C with 15 ng full-length ΔTM gB and gB ectodomains encoding five genotypes in 0.1M Carbonate Buffer. The coating plates were washed once with the plate washer after overnight incubation and blocked with assay diluent (1X PBS containing 4% whey, 15% normal goat serum, and 0.5% Tween 20) at room temperature for 1 hour. Rabbit plasma was 3-fold serial diluted from 1:100 and then added to the plate for 1 hour incubation at room temperature. Cytomegalovirus immune globulin intravenous CytoGam (CSL Behring Healthcare)^83^ was 4-fold serial diluted from 1:1000 and included as assay positive control and plate-to-plate control. The bound rabbit plasma IgG was detected with a horseradish peroxidase (HRP)-conjugated polyclonal mouse anti-rabbit IgG (Southern Biotech) and the CytoGam was detected with an HRP-conjugated polyclonal goat anti-human IgG (Southern Biotech) by 1 hour incubation at room temperature. The plate was later developed with the SureBlue Reserve tetramethylbenzidine (TMB) peroxidase substrate (KPL) and read at 405nm. The 50% effective dose end dilution (ED50) was calculated as the plasma dilution that resulted in a 50% reduction of the IgG binding, determined by the method of Reed and Muench ^84^. The mAb binding domain specificity was determined following the similar procedure described above. 384-well clear-bottom ELISA plates (Corning) were coated with 150 ng gB AD-1 domain (MyBiosource), AD-2 domain site 1 (Life Technologies Corporation), Domain I, Domain II, Domain I+II in 0.1M Carbonate Buffer. The coating plates were washed once with the plate washer after overnight incubation and blocked with assay diluent at room temperature for 1 hour. 2M7 mAb was 4-fold serial diluted from 5 µg/ml and then added to the plate for 2-hour incubation at room temperature. CytoGam (CSL Behring Healthcare), SM-5, SM-10, and TRL-345 was included as assay positive control (Sup Fig. 4A-F). CytoGam was 4-fold serial diluted from 1:1000, while SM-5, SM-10, and TRL-345 was 4-fold serial diluted from 5 µg/ml. 2M7 mAb isolated from rabbit PBMCs was detected with an HRP-conjugated polyclonal mouse anti-rabbit IgG (Southern Biotech) and all the positive controls were detected with an HRP-conjugated polyclonal goat anti-human IgG (Southern Biotech) by 1 hour incubation at room temperature. The plate was later developed with the SureBlue Reserve tetramethylbenzidine (TMB) peroxidase substrate (KPL) and read at 405nm. The 50% effective dose end dilution (ED50) was calculated as the plasma dilution that resulted in a 50% reduction of the IgG binding, determined by the method of Reed and Muench ^84^.

### gB transfected cell binding assay

HEK293T cells (ATCC) were cultured overnight to reach ∼50% confluency in a T75 flask. After overnight incubation, the cells were transfected with 2000ng gB-T2A-GFP plasmids encoding five gB genotypes, respectively, using Effectene Transfection Reagent Kit (Qiagen). T2A peptide, composed of 18-22 amino acids, was found in foot-and-mouth disease virus and commonly known for its self-cleavage ability ^85, 86^. Benefit from the self-cleavage ability of T2A peptide, the GFP structure would not modify the gB structure on the cell surface after transfection. The GFP expression was measured to determine the transfection efficiency of gB-T2A-GFP plasmids. Transfected cells were incubated at 37C, 5% CO2 for 48 hours and then detached with 1X PBS. Cells were re-suspended in DMEM complete media and counted using Countess Automated Cell Counter (Invitrogen). 200,000 cells were plated in a 96-well round bottom plate and blocked with 1:1000 Human TruStain FcX™ (Biolegend) for 10 minutes. After blocking, the cells were washed with 1X PBS once and centrifuged at 1200×g for 5 min to aspirate the washing buffer. Rabbit plasmawas diluted 1:100 and CytoGam was diluted 1:5000 in DMEM complete media, included as positive control and plate-to-plate control. After rabbit plasmaincubation with transfected cells at 37C for 2 hours, cells were washed with 1X PBS+1% FBS and centrifuged at 1200×g for 5 min twice. Dead HEK293T cells were prepared by heating at 95C for 5 minutes as a dead cell control for the following live/dead staining procedure.

The stained cells were washed with 1X PBS+1% FBS and centrifuged at 1200×g for 5 min twice and stained with the 1:1000 Far Red live/dead staining (Invitrogen) at room temperature for 20 minutes. After washing with 1X PBS+1% FBS and centrifuged at 1200×g for 5 min twice, cells were then incubated with PE-conjugated secondary antibody at 4C for 25 minutes. The secondary antibody for the rabbit plasmais PE-conjugated polyclonal mouse anti-rabbit IgG (Southern Biotech), and that for the CytoGam is mouse anti-human IgG (Southern Biotech). The stained cells were washed with 1X PBS+1% FBS twice and fixed with 1X PBS + 4% formalin at room temperature for 15 minutes. After fixation, the fixed cells were washed with 1X PBS+1% FBS twice and resuspended in 100µl PBS for the acquisition. Events were acquired on an LSR-II flow cytometer (BD Biosciences) using the HTS (high throughput screening) cassette. Data was analyzed with Flowjo software (Tree Star, Inc.). GFP expression was measured as the transfection efficiency. As the (**Sup Fig. 1A)** shows, the average transfection efficiency of gB-T2A-GFP plasmids is between 34-49% depends on the gB genotype (gB1: 39%; gB2: 34%; gB-3: 42%; gB-4: 44%; gB-5: 49%). The IgG binding to cell-associated gB was measured by the % of GFP+ PE+ population. Non-specific binding of PE-conjugated mouse anti-rabbit IgG Fc and mouse anti-human IgG Fc was corrected in the analysis. Due to the variations among the transfection efficiency of gB-T2A-GFP plasmids, the IgG binding of week 6, 10, and 30 samples were normalized by GFP expression MFI of week 0 for comparison.

### Glycoprotein B peptide microarray

The rabbit plasma IgG binding to linear 15-mer gB peptides was assessed as previously described ^5, 51^. We designed the 15-mer peptides, overlapping by 10 residues, that covered the amino acids 1-77 of the gB open reading frame from Towne (gB-1), AD169r (gB-2), and Toledo (gB-3) strain since the most variable region lies in the codon 26-70 ^20–22^. A total of 27 peptides were printed in triplicate on the PepStar multi-well array (JPT Peptide). Following the protocol from JPT Peptide, the peptide microarray slide was fixed in the 96-well chamber cassette. The blocking buffer is 3% BSA (Bovine serum albumin) in 1X Tris-buffered saline (TBS)+0.1%Tween20 buffer, and the wash buffer is 1X TBS+0.1%Tween20. Rabbit plasma was centrifuged at 45,000 rpm for 5 min and diluted 1:250 in the blocking buffer. After adding the diluted plasma, the slide was wrapped in aluminum foil and placed on rotator for 1-hour incubation at 150 rpm, 30C. The slide was then washed with 150ul washing buffer four times and incubated with 1:500 Alexa Fluor® 647 AffiniPure Goat Anti-Rabbit IgG (H+L) (Jackson Immuno Research) for 1-hour incubation at 150 rpm, 30C. The stained slide was washed with 150ul washing buffer four times again and further washed with 150ul DI water twice before drying by centrifuge spin at 1,350 rpm for 5 min. The dried microarray slide was scanned at 635nm by Genepix 4100A and analyzed by GenePix Pro 6.1 software (Molecular Devices). The plasma binding to linear 15-mer gB peptides was calculated by (feature intensity-background intensity) at 635nm, and the median peptide binding of each of the triplicates was reported. The cutoff for positive peptide binding was defined by the average binding of the secondary non-specific antibody binding, Alexa Fluor® 647 AffiniPure Goat Anti-Rabbit IgG (H+L), plus two standard deviations.

### Neutralization

The fibroblast neutralization titer was measured on HFF-1 cells (ATCC) and the epithelial neutralization titer was detected on ARPE-19 cells (ATCC). 5000 cells/well HFF-1 cells or ARPE-19 cells were resuspended in complete DMEM media+20% FBS or complete DMEM media+10% FBS, respectively. HFF-1 cells or ARPE cells were later plated in 384-well black and clear-bottom plates (Corning) overnight at 37C. Rabbit plasmawas heat-inactivated at 65C for 1 hour to inactivate the proteins that impact the neutralization activity (e.g., complement). Next, rabbit plasma was 3-fold serial diluted from 1:8 and CytoGam was 3-fold serial diluted from 1:80, included as assay positive control and plate-to-plate control. The diluted rabbit plasma was incubated with Towne, AD169r, or Toledo virus (a kind gift from Dr. Ravit Boger), respectively, at 37C, 5% CO2 for 1 hour. To assess the rabbit plasma neutralization antibody response with the effect of complement, purified rabbit complement (Cedarlane Laboratories) was mixed in the complete DMEM media+20% FBS at a ratio of 1:4. After 1-hour incubation, the mixture of rabbit plasmaand HCMV was added in duplicate to wells containing HFF-1 or ARPE-19 cells for a further incubation. HFF-1 cells were incubated at 37C, 5% CO2 for another 20-24 hours, while ARPE-19 cells were further incubated at 37C for 44-48 hours. Infected cells were then fixed with 1X PBS + 4% formalin at room temperature for 15 minutes before staining. The staining buffer is 1X Dulbecco’s PBS (DPBS) + 1% FBS + 0.3% Triton X-100. Fixed cells were stained with 1:1000 mouse anti-HCMV IE-1 monoclonal antibody (MAB810; Millipore) for 1 hour and then stained with 1:1000 goat anti-mouse IgG-AF488 (Millipore) for another 1 hour at room temperature. The nuclear stain was later performed by incubating DAPI (4=,6-diamidino-2-phenylindole) nuclear stain for 10 minutes at room temperature. After staining, the cells were resuspended in 1X PBS before detection. The total cells and AF488+ cells were counted on a Cellomics ArrayScan reader (Thermo Fisher Scientific) or ImageXpress Pico Automated Cell Imaging System (Molecular Devices). The % infection rate was determined by the AF488+ cells/total cells. Neutralization titers (ID50) were calculated based on 50% reduction of the % infected cells via the method of Reed and Muench ^84^.

### ADCP of whole HCMV virions

Towne, AD169, and Toledo virus was conjugated with DMSO-dissolved AF647–N-hydroxysuccinimide ester (Invitrogen) with constant agitation at room temperature for 1 hour. The conjugation reaction was quenched with 1 M Tris-HCl, pH 8.0. In a 96-well round-shaped plate, rabbit plasma was diluted 1:30 in 1X PBS+ 1% FBS. CytoGam (CSL Behring Healthcare) was 3-fold serial diluted from 1:30 and as positive control and plate-to-plate control, while PBS was included as negative control. The diluted sera and controls were incubated with viruses at 37C for 2 hours. After 2-hour incubation, 250,000 cells/ml THP-1 cells (ATCC) were resuspended in RPMI+10% FBS media and 200µl THP-1 cells were added to the sera-virus mixture. The 96-well plate was centrifuged at 1,200 xg, 4C for 1 hour to allow spinoculation and then incubated at 37C, 5% CO2 for an additional 1 hour. The cells were spun down at 1,200 xg for 5 minutes and washed with 1X PBS+ 1% FBS once before staining with Aqua Live/Dead stain (Invitrogen) at room temperature for 25 minutes. The stained cells were washed with 1X PBS+1% FBS twice and fixed with 1X PBS + 4% formalin at room temperature for 15 minutes. After fixation, the fixed cells were washed with 1X PBS+1% FBS twice and resuspended in 100 µl PBS for the acquisition. Events were acquired on an LSR-II flow cytometer (BD Biosciences) using the HTS (high throughput screening) cassette. Data were analyzed with Flowjo software (Tree Star, Inc.). The ADCP activity was determined by the AF647+ cells from the live cell population using the PBS negative control as the threshold **(Sup Fig. 3A-B)**.

### Monoclonal antibody screening and isolation

Rabbit memory B cells culture followed by mAb screening and cloning was previously reported ^45, 87^. The week 10 rabbit PBMCs from the monovalent and pentavalent vaccine groups (around 1E+07 cells per sample, 1 animal from each group) were mixed to ensure the cell number is enough for the following cell culture. A total of 600 IgG+ memory B cells were incubated and enriched with HCMV AD169r strain, then seeded into 384-well plates, at a density of one cell/per well with pre-coated EL4-B5 feeder cells (Kerafast). The feeder cells were irradiated with 50 Gy in a gamma radiation chamber before use. The single memory B cells were cultured in complete RPMI 1640 medium supplemented with IL-2 (10 U/ml, R&D Systems), IL-21 (10 U/ml, R&D Systems), at 37C, 5% CO2, 93% humidity for 14 days. The B-cell culture supernatants were collected. These supernatants were tested in post-fusion full-length gB binding ELISA and AD169r neutralization assay to screen the positive cell candidates. The total RNA from the cell candidates was isolated with RNeasy Micro Kit (Qiagen) and converted to cDNA using iScript™ cDNA Synthesis Kit (Bio-Rad). The IgG heavy and light chain genes were amplified by PCR as previously described^45^. The plasmids encoding the heavy and light chain genes were later transiently transfected in Expi293F cells (Thermo Fisher Scientific), and the recombinant mAbs were purified by Protein A/G affinity chromatography as reported^45^.

### gB-specific T cell IFN-γ ELISpot assay

Rabbit IFN-γ ELISpot BASIC kit (MabTech) was used to determine IFN-γ production from rabbit PBMCs and splenocytes. Following the protocol from MabTech, 96-well MultiScreenHTS IP Filter Plate (Millipore) was activated with 35% ethanol and coated with 15 µg/ml MT327 coating antibody overnight at 4C. On the following day, the 96-well plate was washed with sterile PBS five times and blocked with RPMI+10% FBS media 30 minutes at room temperature. Frozen rabbit primary PBMCs and splenocytes were thawed in RPMI+10% FBS media and counted by Countess™ 3 Automated Cell Counter (Invitrogen). 200,000 primary rabbit cells were plated in each well and stimulated with PepMix™ HCMVA UL55 peptides (JPT Peptide Technologies) overnight at 37C, 5% CO2. Cells were also stimulated with 0.4µg/ml PMA (MilliporeSigma) and 4 µg/ml ionomycin (Sigma-Aldrich) for positive control and DMSO (Sigma-Aldrich) or RPMI+10% FBS media only for negative control. After 20-hour incubation, cells were removed, and the plate was washed with sterile PBS five times before staining. The staining buffer is 1X DPBS + 0.5% FBS. The secreted cytokine was incubated with 0.1µg/ml MT318-biotin detection antibody in staining buffer for 2 hours at room temperature, followed by five-time sterile PBS wash. The plate was later incubated with 1:1000 Streptavidin-ALP conjugate in staining buffer for 1 hour at room temperature, followed by five-time sterile PBS wash. Alkaline phosphate conjugate substrate kit (Bio-Rad) was applied for substrate color development. The plate was developed with color development solution for 25 minutes at room temperature and the reaction was stopped by incubating with 0.5% Tween-20 in 1X DPBS for 10 minutes. After removing the color development solution, the plate was washed with tap water four times and dried overnight before reading using ImmunoSpot® S6 Ultimate M2 Analyzer (Cellular Technology Limited). The IFN-γ-positive spots were imaged and counted using ImmunoSpot® Software (Cellular Technology Limited).

### Statistical analysis

In the longitudinal analysis of Full-Length gB IgG Elisa we utilized a non-parametric repeated measures test due to the small sample with permutations used due to a singular covariance matrix ^88, 89^. For the TCB and ectodomain analysis, a multivariate Kruskal-Wallis test was used with 100000 permutations to conduct inference using the coin package in R ^90^. For the neutralizing and ADCP analysis, Kruskal-Wallis was again used-with permutations due to the small sample size at each time point and virus ^90^. For all analyses, multiple testing correction was done used FDR ^91^ with significance at a FDR adjusted p-value less than 0.1 and an unadjusted p-value less than 0.05. This was with the exception of when only adjusted for two tests in which a Bonferroni correction was used. Any post-hoc analyses (within specific virus), multiple testing correction was done using the FWER controlling Holm procedure^88^. Secondary tests between individual vaccines (Monovalent vs Bivalent/Pentavalent combined for example) were done using the Wilcoxon Rank Sum test in the coin package ^90^. All analyses were done within R ^92^. For the T cell response analysis, we performed Kruskal-Wallis as above with multiple testing correction via Bonferroni.

## Acknowledgement

We thank Dr. Herman Staats at Duke University for providing us New Zealand White rabbits and the animal handling tips for our study. We also thank Duke Animal Handling Facility for taking care of animals and assistance with immunizations and sample collections. As for the experimental reagents, we appreciate Sanofi Inc for providing us post-fusion full-length gB. Also, we appreciate Dr. Ravit Boger from Johns Hopkins University for providing us Toledo virus. We wish to acknowledge following funding sources for supporting this project: 1) Biostatistics, Epidemiology and Research Design (BERD) Methods Core funded through Grant UL1TR002553 from the National Center for Advancing Translational Sciences (NCATS), a component of the NIH, to CC. 2) 5P01AI129859 from the National Institute of Allergy and Infectious Diseases (NIAID), a component of the NIH, to SRP. 3) Cancer Prevention and Research Institute of Texas (RP150551 and RP190561) and the Welch Foundation (AU-0042-20030616) to ZA. 4) Medearis CMV Scholar Award from Duke University Medical Center to HYW.

## Figure Citations

**Sup Fig 1**. ^51, 60, 93, 94^

---

**Sup Figure 1.**
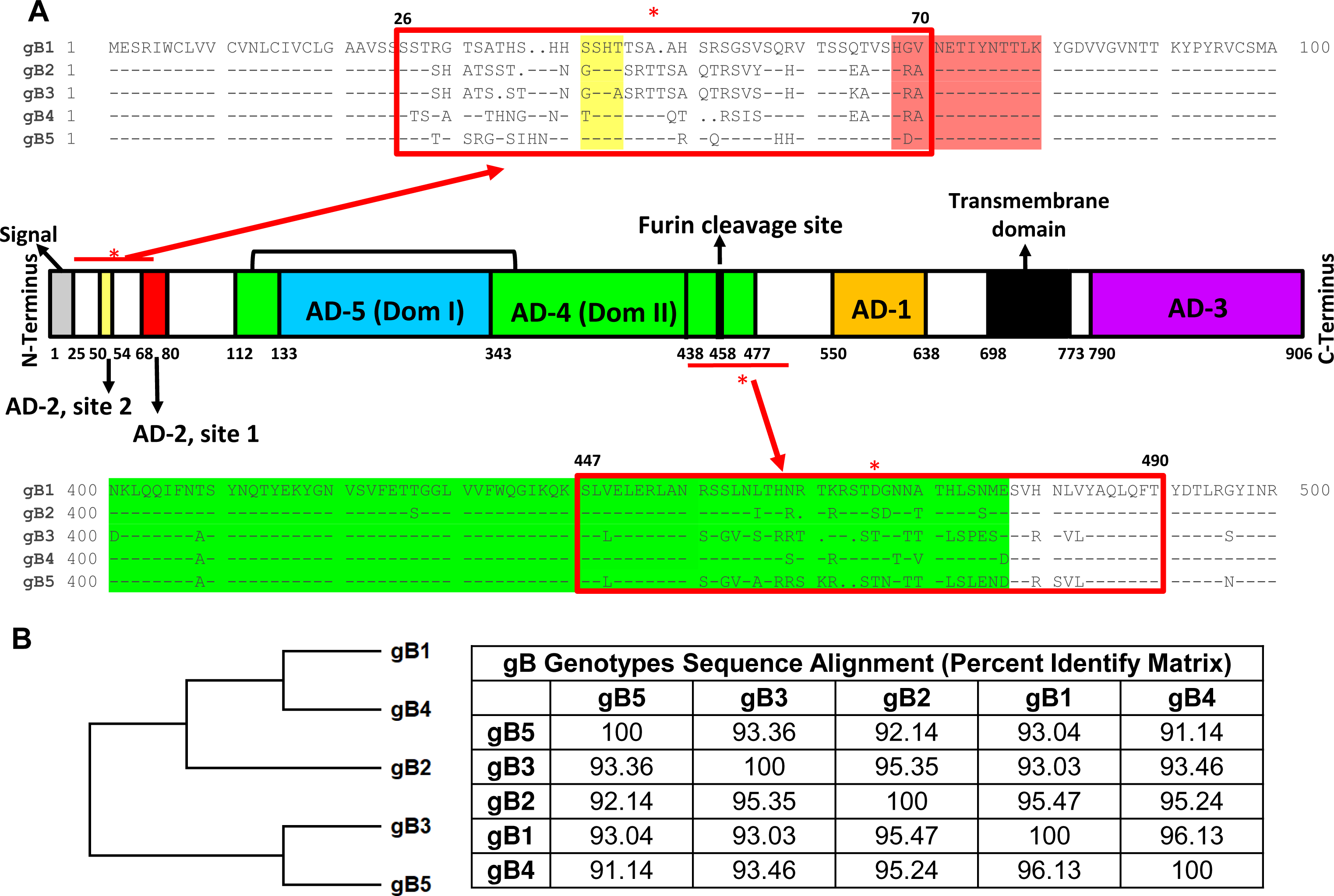
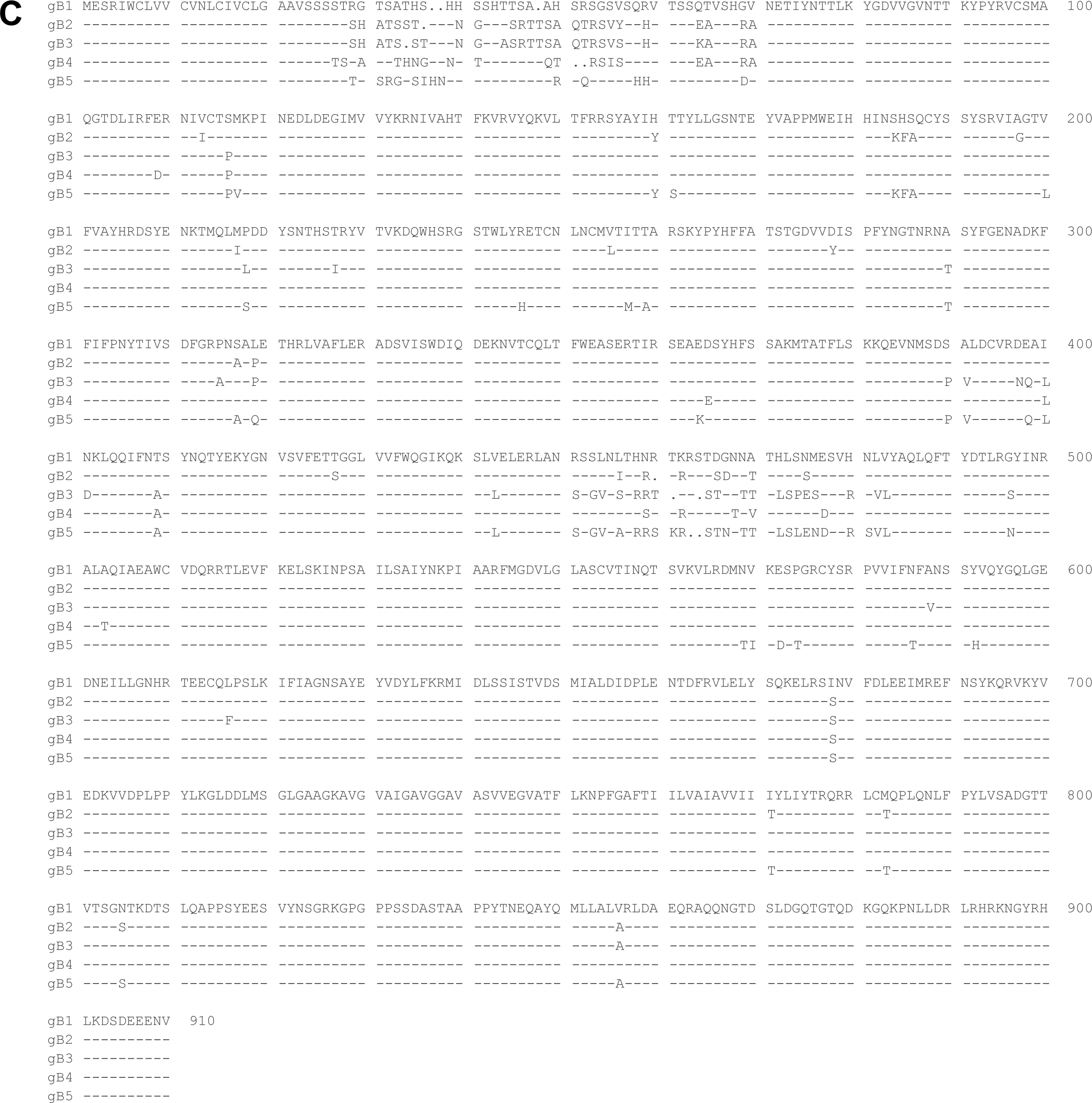
Schematic representation of HCMV gB sequence with antigenic domains and the most variable regions among 5 gB genotypes. (A) The full opening reading frame of HCMV gB is shown from the N-terminus (left) to the C-terminus (right). The gB sequence consists of ectodomain, transmembrane domain (black), and the cytosolic domain. The AD-1 (orange), AD-2 site 1 (red), AD-2 site 2 (yellow), AD-4 (also known as Dom II, green), and AD-5 (also known as Dom I, blue) are in ectodomain, while AD-3 (purple) belongs to cytosolic domain. This figure was adapted from Burke et al. (60), Chandramouli et al. (93), and Nelson et al. (51). The most variable regions defining the five gB genotypes are codons 26-70 and codons 441-490. These regions were aligned using Clustal W and Clustal X version 2.0 (94) and presented by SeqPublish. The five gB sequences included in the vaccine design are: gB-1 (C327A strain, M60929.1), gB-2 (C336A strain, M60931.1), gB-3 (BE/37/2011 strain, KP745723.1), gB-4 (C194A strain, M60926.2), and gB-5 (17_saliva_7-24-2003 isolate, MK157431.1). (B) Phylogenetic tree of full gB open reading frame encoding 5 gB genotypes. The phylogenetic analysis was performed following the bootstrap method with 100 replications by Molecular Evolutionary Genetics Analysis version 11 (Tamura 2011). Clades representing 5 individual gB genotypes are color-coded : gB-1, black circle; gB-2, red square; gB-3, green upward triangle; gB-4, blue diamond; gB-5, orange downward triangle. The precent identity matrix comparing five gB genotypes was generated using Clustal W and Clustal X version 2.0 (94). (C) The complete sequence alignment of the five HCMV gB genotypes using Clustal W and Clustal X version 2.0 (94) and presented by SeqPublish. The five gB sequences included in the vaccine design are: gB-1 (C327A strain, M60929.1), gB-2 (C336A strain, M60931.1), gB-3 (BE/37/2011 strain, KP745723.1), gB-4 (C194A strain, M60926.2), and gB-5 (17_saliva_7-24-2003 isolate, MK157431.1).

**Sup Table 1.**
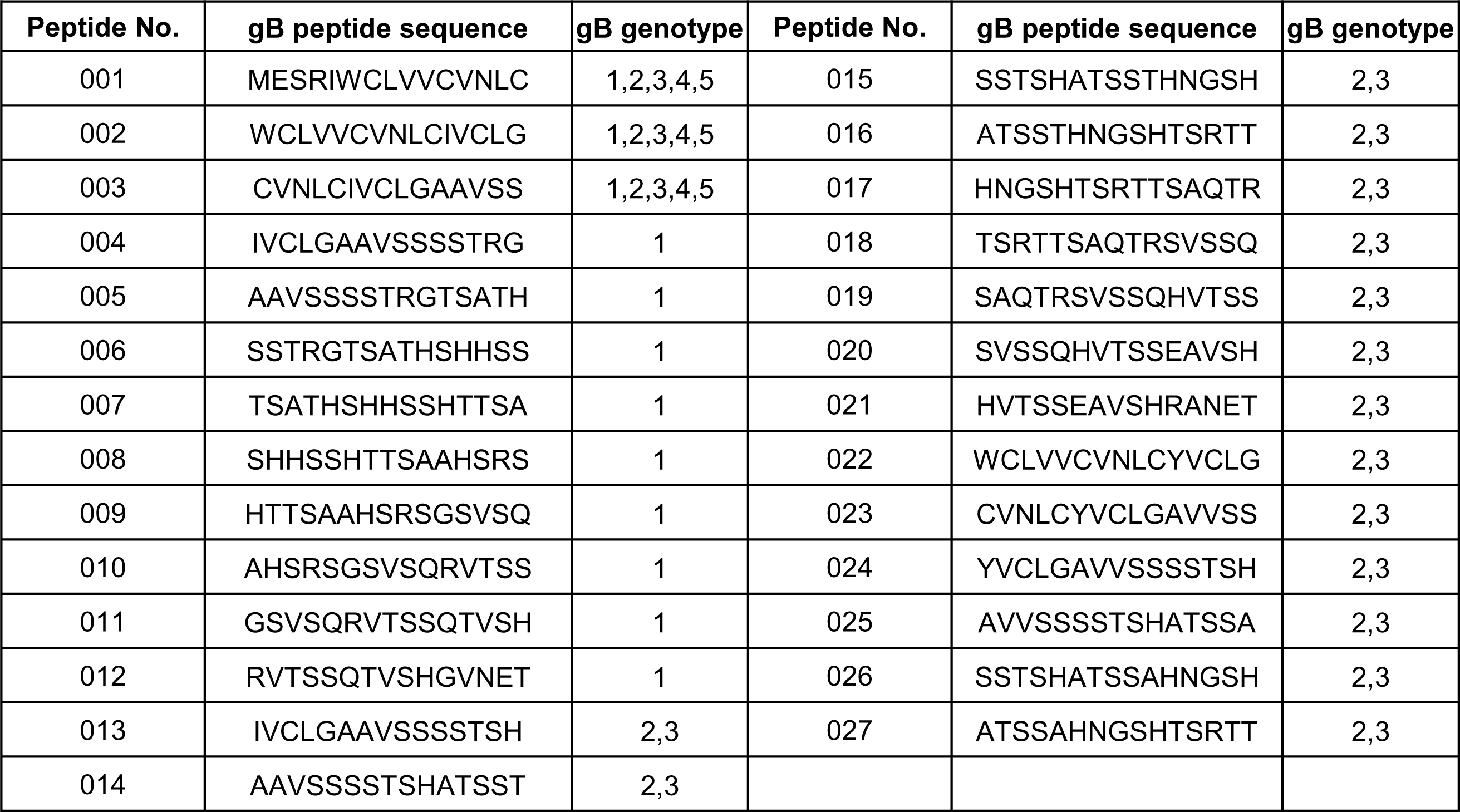
Linear overlapping 15-mer gB peptide sequence.

**Sup Figure 2.**
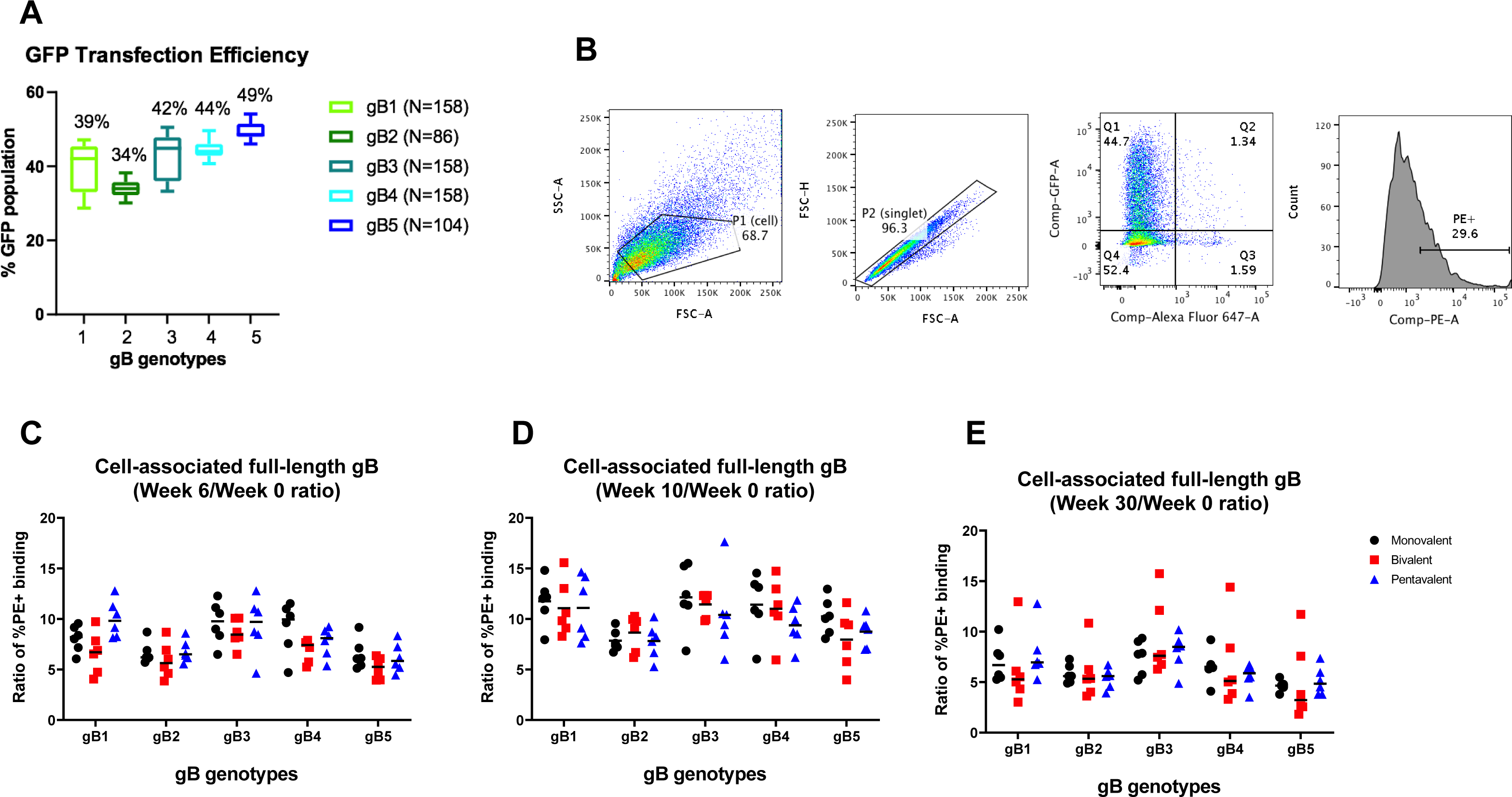
Monovalent, bivalent, and pentavalent gB mRNA-LNP vaccinated rabbit plasma IgG elicited a similar binding breadth to cell-associated gB of 5 genotypes. (A) The transfection efficiency of gB-T2A-GFP plasmids encoding 5 genotypes, respectively, was determined by %GFP expression. The %GFP expression was shown by box and whisker plot, with the mean value labeled on top. These values were color-coded based on the gB genotypes: gB-1, lawn green; gB-2, Forest green; gB-3, Paris green; gB-4, sky blue; gB-5, blue. (B) Gating strategy of gB transfected cell binding assay. The 293T cells were determined by SSC-A and FSC-A, designated as P1 population. The P2 singlet population was later obtained by gating the P1 population through FSC-A and FSC-H. Next, the live GFP+ P2 population was shown in Q1, and the %PE+ population was compared from the Q1 population. (C-E) Rabbit plasma IgG binding breadth to full-length gB encoding 5 genotypes at week 6 (C), week 10 (D), and week 30 (E) by gB-transfected cell binding assay. Data points are shown as the IgG binding response normalized to that of the week 0 samples to eliminate the discrepancies among the transfection efficiency. Each data point represents the normalized IgG binding response of one individual animal, with the median response labeled by a black line. Black circles: rabbits immunized with monovalent vaccine; red squares: those immunized with bivalent vaccine; blue triangles: those immunized with pentavalent vaccine.

**Sup Figure 3.**
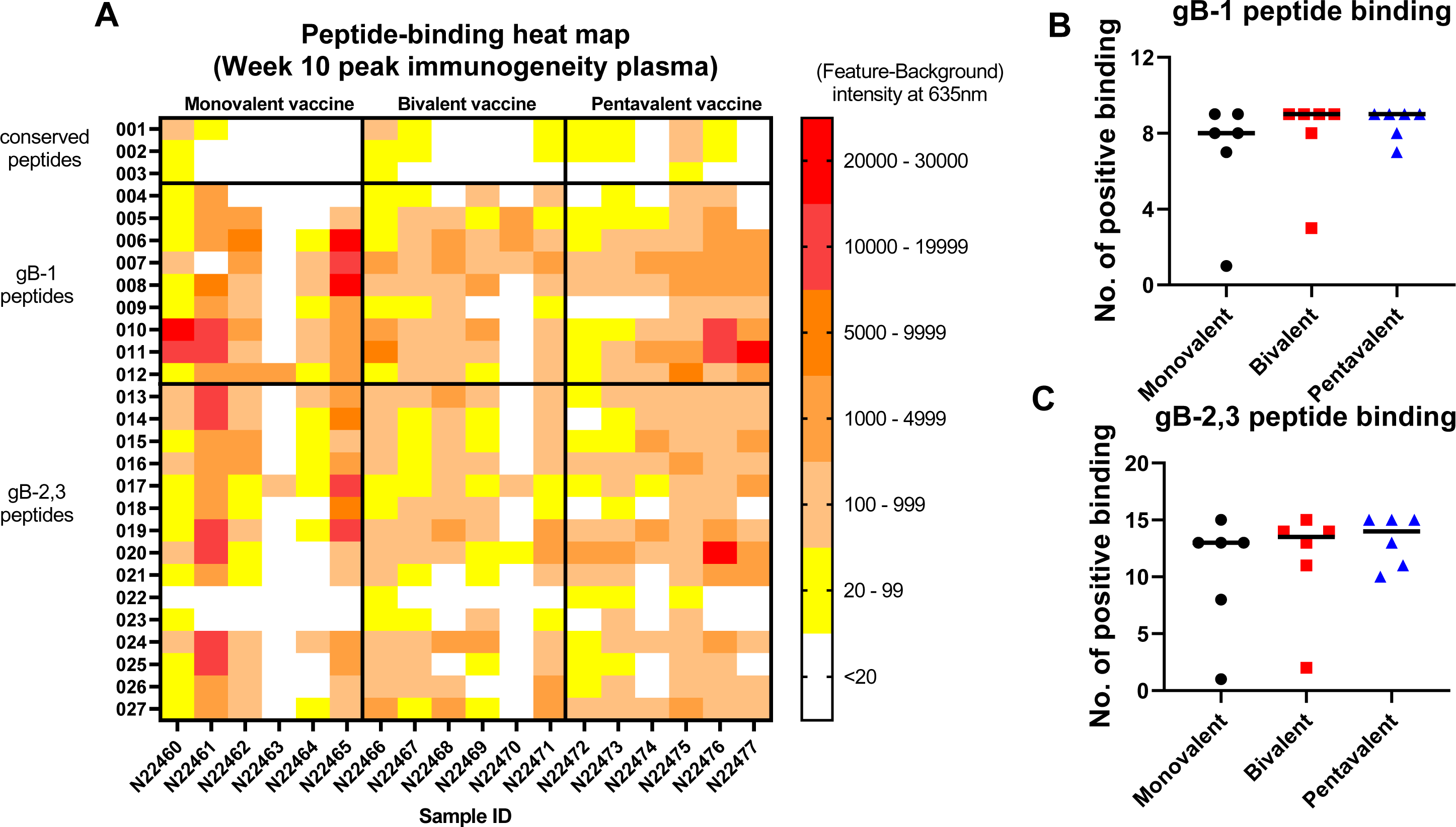
Monovalent, bivalent, and pentavalent gB mRNA-LNP vaccinated rabbit plasma IgG at week 10 did not show a statistical different binding pattern to linear gB peptides of gB-1 or gB-2 and 3 genotypes. (A) The breadth of rabbit plasma IgG binding to 15-mer linear gB peptides covering the amino acid codons 1-77 of gB-1 or gB2/3 genotypes (27 unique peptides) at peak immunogenicity (week 10), measured by peptide microarray. Each row indicates a linear gB peptide: 001-003, conserved peptides; 004-012, gB-1 peptides; 013-027, gB2/3 peptides. Each column is one animal from each vaccine group. The plasma IgG binding to linear 15-mer gB peptides was calculated by (feature-background) intensity at 635nm, and the average peptide median binding (feature-background) intensity at 635nm of each of the triplicates was reported. (B-C) The positive binding MFI cutoff was defined by the average binding of the secondary non-specific antibody binding plus two standard deviations. Each data point represents the number of peptides bound by plasma IgG from one individual animal, with the median response labeled by a black line. Black circles: rabbits immunized with monovalent vaccine; red squares: those immunized with bivalent vaccine; blue triangles: those immunized with pentavalent vaccine.

**Sup Figure 4.**
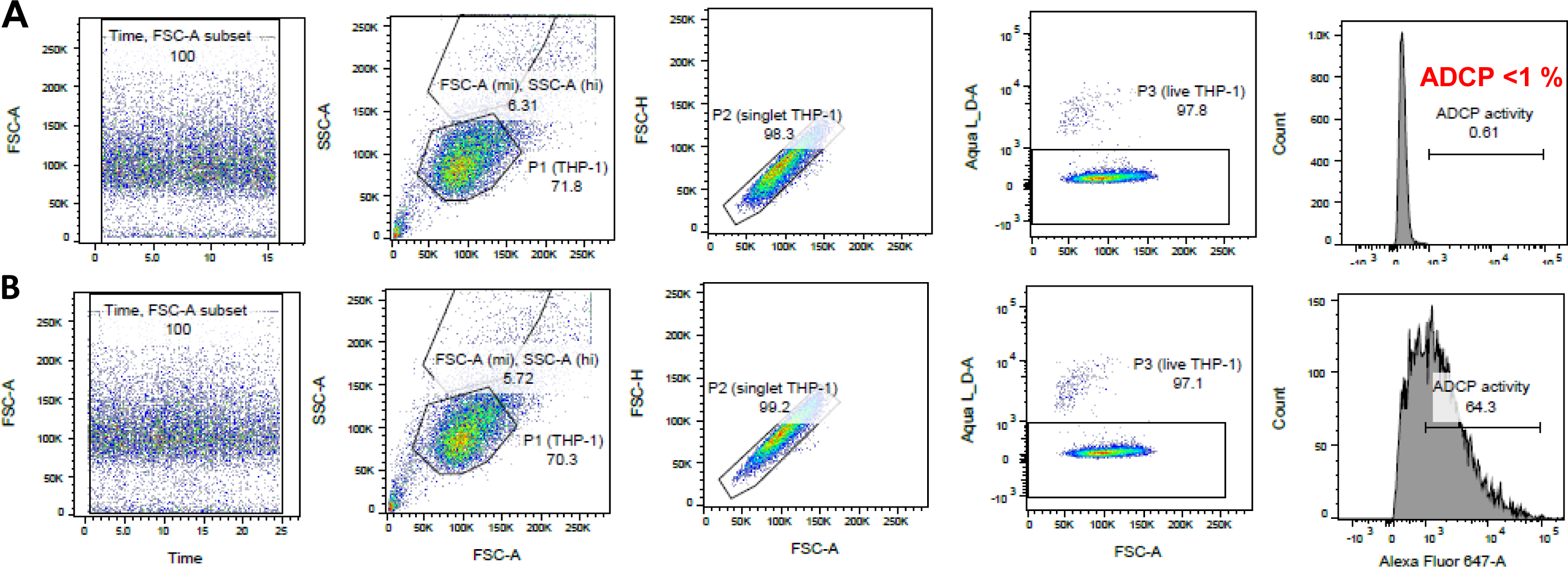
Gating strategy of ADCP for negative control (PBS) and positive control (CytoGam). (A-B) The THP-1 cells were first gated for acquisition time and the cells were determined by SSC-A and FSC-A, designated as P1 population. The P2 singlet THP-1 population was later obtained by gating the P1 population through FSC-A and FSC-H. Next, the live THP-1 cell population was by aqua L/D stain as P3, and the % ADCP activity was determined by the AF647+ cells from the live cell population using the PBS negative control as the threshold. ADCP activity from PBS negative control (A) and CytoGam positive control (B).

**Sup Figure 5.**
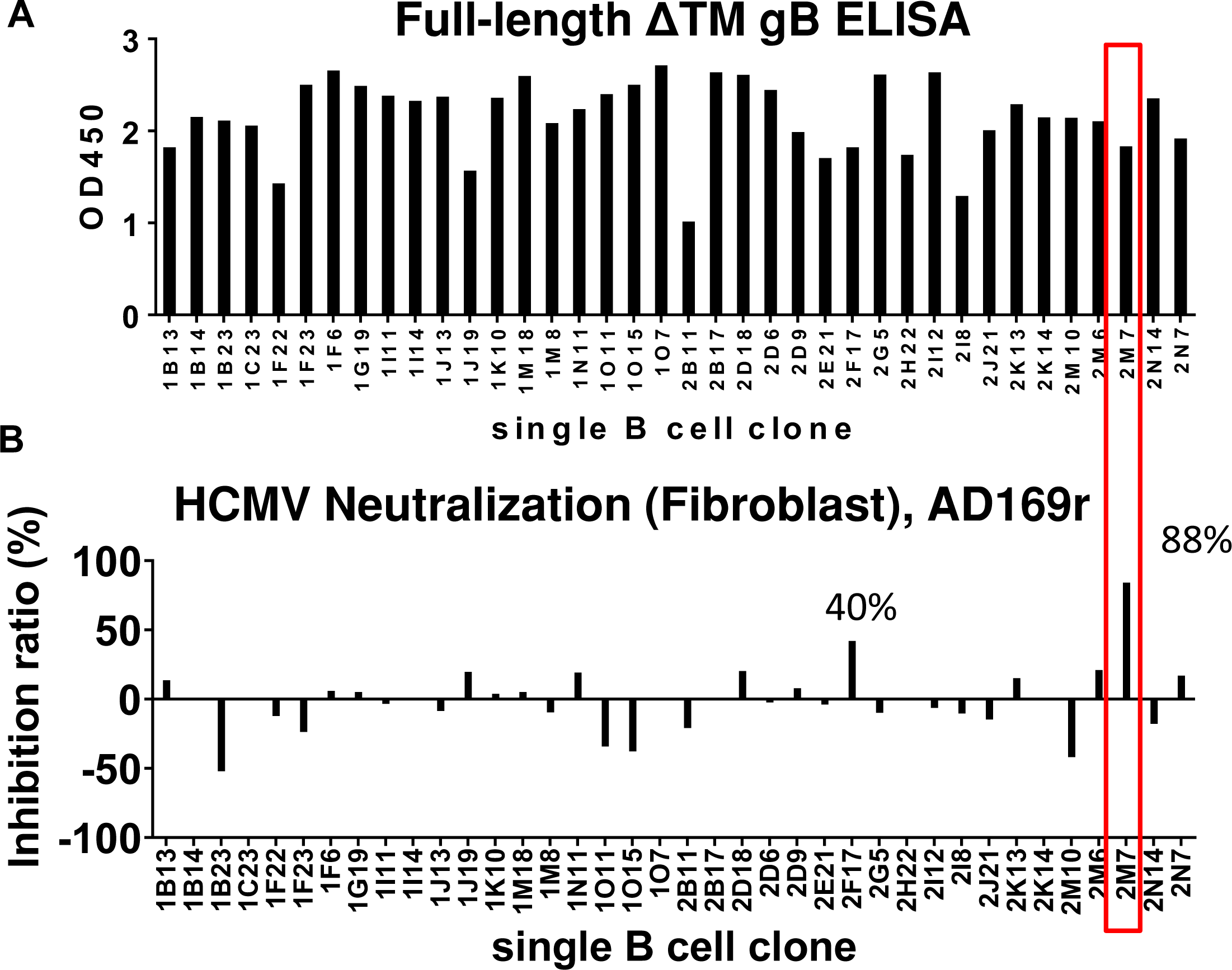
Monoclonal antibody screening from memory B cell clones. (A) Among 600 single memory B cell clones, the supernatant (1:20 dilution) of 38 clones demonstrated a binding response to post-fusion full-length gB by ELISA. (B) The fibroblast neutralization potency of the supernatants from the 38 gB-binding clones (1:20 dilution) was examined by neutralization assay against HCMV AD169r strain. Among 38 mAbs, only 2M7 showed the strongest neutralization potency.

**Sup Figure 6.**
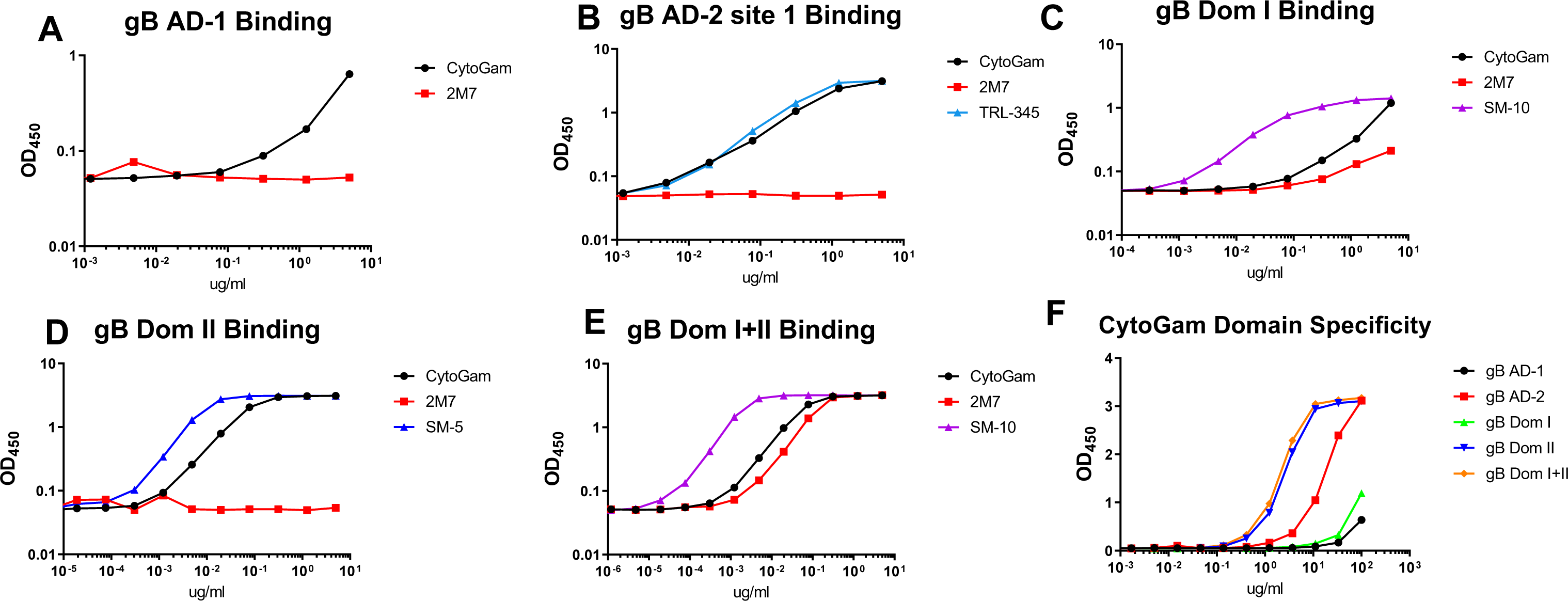
Positive controls of gB domain specificity confirmed 2M7 only showed binding to gB Dom I+II. (A-E) The gB domain specificity of 2M7 was determined by comparing binding to gB AD-1 (A), AD-2 (B), Domain I (C), Domain II (D), and domain I+II (E). The positive controls for each antigen are CytoGam (A-F, black circle), TRL-345 (B, light blue triangle), SM-10 (C, E, purple triangle), and SM-5 (D, blue triangle). (F) The gB domain specificity of CytoGam, measured by ELISA. CytoGam binding to gB domains is color-coded with symbols: black circle, AD-1; red square, AD-2 site 1; green upward triangle, Domain I; blue downward triangle, Domain II; orange diamond, Domain I+II.

**Sup Figure 7.**
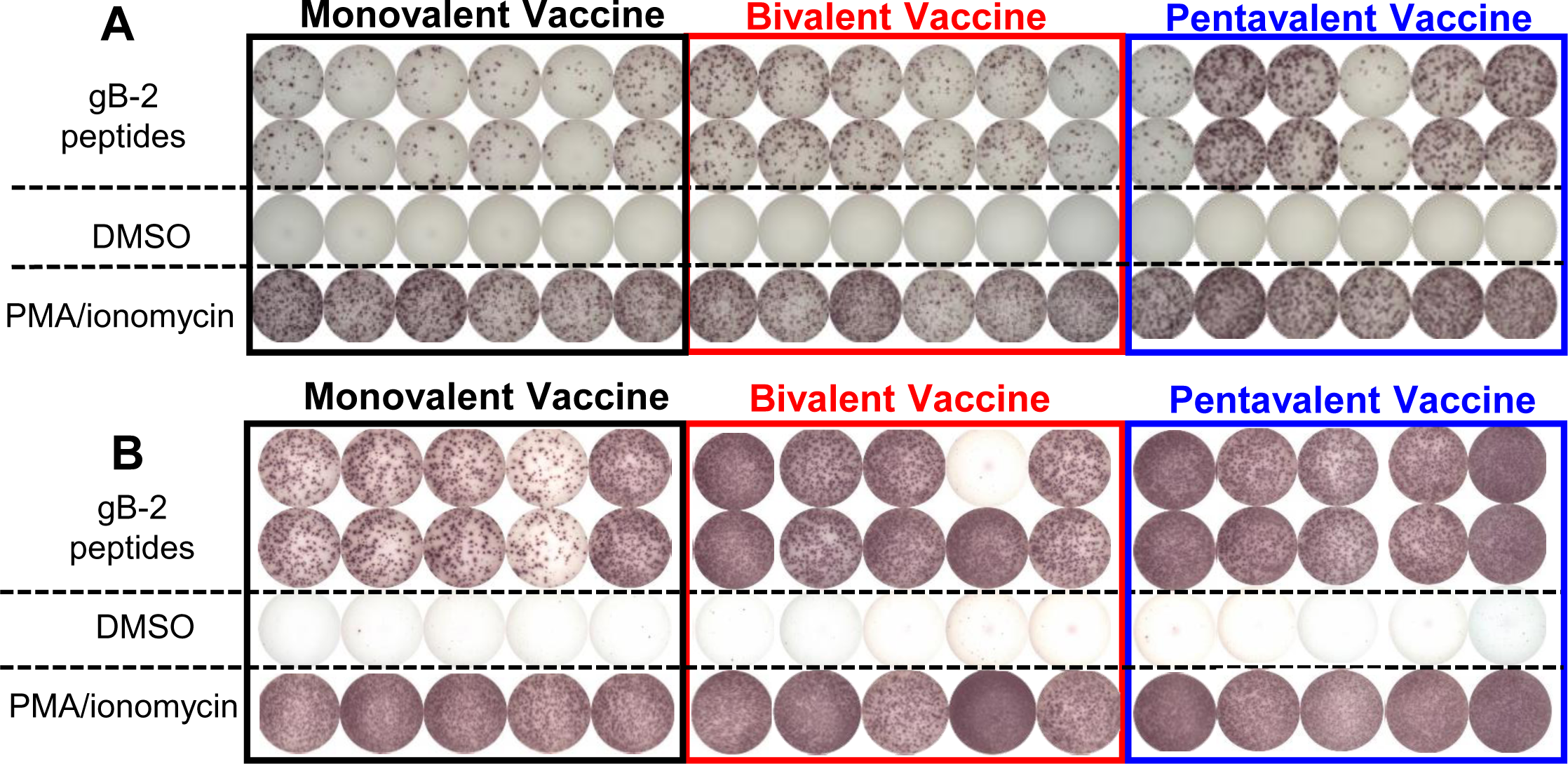
Representative wells of IFN-r+ cells simulated with gB peptides, DMSO, or PMA/ionomycin from rabbit PBMCs at week 6 and splenocytes at week 30. gB-2-specific IFN-r+ cells from rabbit PBMCs or splenocytes were measured by ELISPOT. gB-2 peptide pool stimulation were performed in duplicate. Black rectangle: rabbits immunized with monovalent vaccine; red rectangle: those immunized with bivalent vaccine; blue rectangle: those immunized with pentavalent vaccine. Representative wells of rabbit PBMCs (A) and splenocytes (B). (B) One well from the bivalent vaccine group incubated with gB-2 peptides did not show any binding and thus was removed for analysis.

